# MSK1 absence hinders BDNF-dependent striatal neurodevelopment and leads to schizophrenia symptoms

**DOI:** 10.1101/2024.01.23.576945

**Authors:** Natalia Varela-Andrés, Alejandro Cebrián-León, Carlos Hernández-del Caño, Inés S. Fernández del Campo, Sandra García-Losada, Noelia Martín-Ávila, Juan Carlos Arévalo, Miguel A. Merchán, Manuel Sánchez-Martín, Rubén Deogracias

## Abstract

Brain-derived neurotrophic factor (BDNF) plays a critical role in postnatal development by modulating the architecture of specific neuronal populations and brain areas. However, the precise molecular program controlling this differential responsiveness to BDNF is still unclear. In the present study, we describe that this program is governed by the restricted expression of the mitogen- and stress-activated protein kinase-1 (MSK1) in GABAergic neurons. Also, we show that while *Msk1* expression declines in cortical interneurons along early postnatal development, its expression in striatal neurons increases until adulthood. Utilizing a novel MSK1 loss-of-function mouse model, we reveal its essential role in postnatal growth of the striatum, as it interacts with and modulates the BDNF-dependent phosphorylation of the methyl-CpG binding protein-2 (MeCP2). Furthermore, these mutant mice exhibit an altered transcription pattern of genes involved in the control of the dopamine and GABAergic signalling pathways. Consequently, MSK1 knockout mice behaviour is markedly altered, showing social dysfunction, altered anxiety- and depressive-like responses unequally manifested in males and females. These results elucidate how disruptions in the BDNF/MSK1 pathway impact GABAergic neurite outgrowth and contribute to behaviours reminiscent of schizophrenia in humans.

## Introduction

During brain development, both extrinsic and intrinsic factors play a crucial role in shaping neural circuits and establishing connectivity patterns between neurons. Neurotrophins, in particular the brain-derived neurotrophic factor (BDNF) (1–5), neurotransmitters such as GABA, glutamate or dopamine (6–10), DNA-binding proteins such as MeCP2 (11–13) or transcription factors like cAMP/Ca2^+^ response element-binding protein (CREB) (14–16) stand out. In combination, these molecules ensure the proper formation and organization of neural circuits and control precise synaptogenesis.

*Bdnf* deletion experiments have revealed that the morphology of GABAergic neurons is much more impacted by BDNF deprivation than is the case for excitatory neurons (17–19). This program allows BDNF to regulate striatal growth and medium spiny neurons (MSNs) arborization (18). Among the different signalling pathways that mediate BDNF-dependent effects, the MAPK pathway plays a role in regulating the branching of MSNs (20). This pathway involves the activation of ribosomal S6 kinase (RSK1, RSK2, RSK3) and mitogen- and stress-activated kinase protein families (MSK1, MSK2) (21–25) in different cell and neuronal types. Both protein families promote gene expression through the activation of transcription factors such as CREB (26, 27).

MSK1 is recruited to immediate-early gene (IEG) regulatory regions (27), where it phosphorylates Histone H3 and CREB (28) to expand the dynamic range of synapses in the hippocampus (29). Studies with adult mice lacking either MSK1 or both MSK1 and MSK2 have shown that MSK1 is involved in the experience-dependent enhancement of mental ability, neuronal function, and cognitive performance (30, 31). Additionally, cocaine administration activates MSK1 in the striatum, while MSK1 lack causes an increased response to low doses of this drug. This suggests the potential role of this kinase in modulating neuronal responses to several stimuli (31, 32). As *Msk1* is highly expressed in striatum and cerebellum in adult mice (33, 34), we hypothesized that it could play an important role during mouse GABAergic development. The synthesis of GABA by the enzymes GAD65 and GAD67 plays a crucial role in the maturation and functionality of inhibitory circuits in the brain (35). The expression of *Gad2* (GAD65) is controlled by BDNF and CREB activation (3).

To study the possible link between MSK1 and GABAergic development, we analysed the expression pattern of *Msk1* at early postnatal ages and found it to be high in both GABAergic cortical interneurons and MSNs. Surprisingly, *Msk1* expression on these populations is both area- and age-specific regulated at least until young-adulthood. In a newly generated *Msk1* knockout murine model (*Msk1^IV^* KO), we observed a significant impact on the growth of the striatum. Also, we found that MSK1 is a critical factor for the arborization of cultured striatal neurons. Furthermore, we observed that MSK1 interacts with MeCP2, controls its BDNF-dependent phosphorylation, and regulates the expression of genes involved in the GABA and dopamine functions in the striatum.

Finally, we observed that the lack of MSK1 alters the innate and specific social responses, anxiety, and depressive behaviours in mice without affecting their locomotor activity. These behavioural features are strongly reminiscent of phenotypes observed in schizophrenia patients.

## Results

### *Msk1* expression changes during early postnatal mouse brain development

To study the expression pattern of *Msk1* during mouse brain development, we analysed both the striatum and the somatosensory cortex between postnatal days P5 to P30 by qPCR and western blot assays (Fig. 1A and B). Msk1 transcript levels are very low in the somatosensory cortex between P5 and P10+, reaching nearly half of the amount of its levels at P30 (Fig. 1A). However, we observed a significant increment of *Msk1* mRNA levels in the striatum during all the analysed time points, with two significant differences between P5 to P10, and between P15 to P30 (Fig. 1A). MSK1 protein levels vary following a similar expression pattern to its transcripts at the same time points in both brain areas (Fig. 1B).

**Figure 1.**
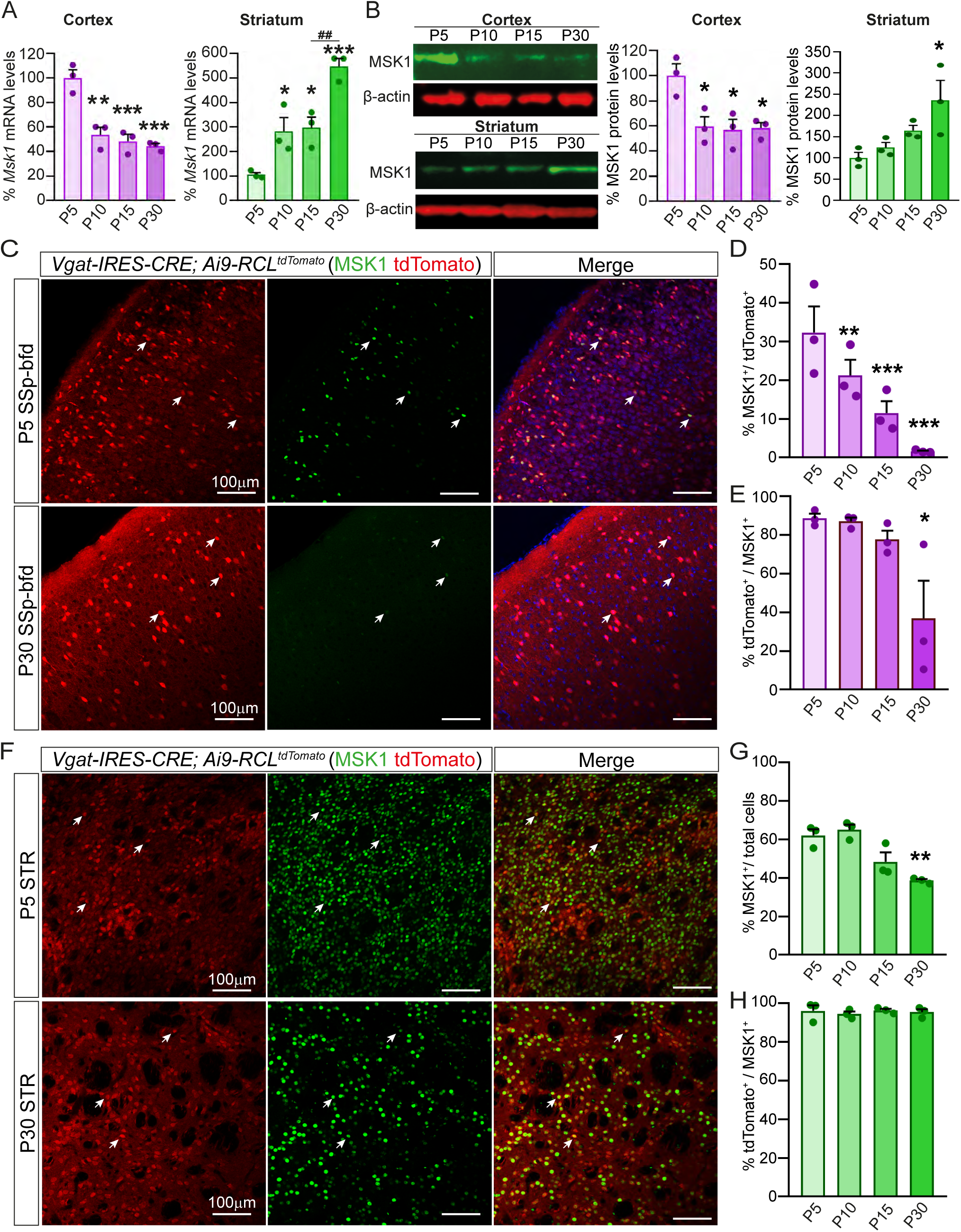
Differential expression of MSK1 during postnatal mouse brain development in cortex and striatum. (A) *Msk1* mRNA expression levels (qPCR) in cortex and striatum at different postnatal ages (P5, P10, P15 and P30). n=3 animal/age group. (B) MSK1 expression levels (WB) in cortex and striatum at different postnatal ages using specific antibodies against the C-terminal domain of MSK1. n=3 animals/age group. (C and F) Representative single plane confocal images of MSK1 expression pattern (green) in the somatosensory cortex and striatum of 5- and 30-day-old *Vgat-IRES-CRE; Ai9-RCL^tdTomato^* mice. Endogenous tdTomato fluorescence (red) reveals GABAergic neurons. Arrows point to cells co-expressing MSK1 and tdTomato. (D and E) Quantification of MSK1^+^/tdTomato^+^ cells and tdTomato^+^/MSK1^+^ cells in the somatosensory cortex at different postnatal ages. n=3 mice/age group, 6-8 images per animal. (G and H) Quantification of MSK1^+^ cells and tdTomato^+^ MSK1^+^ cells in the striatum at different postnatal ages. n=3 mice, 6-8 images per animal; Scale bars = 100µm. Results are the mean ± SEM. Statistical analysis was performed using one-way ANOVA; \**P*<0.05, \*\**P*<0.01, ^##^*P*<0.01, \*\*\**P*<0.001.

### MSK1 is mostly restricted to GABAergic neurons

The changes observed on MSK1 protein levels during brain development might not be uniform throughout all the cells in the somatosensory cortex and the striatum. In the adult mouse brain *Msk1* is expressed by dopamine- and cAMP-regulated phosphoprotein, 32 kDa (DARPP-32) positive MSNs (36) and by few neurons in the cortex (33). We hypothesized that MSK1 could be mostly restricted to cortical GABAergic interneurons, while in the striatum MSK1 could be located in the GABAergic MSNs. To assess this, we used the *Vgat-IRES-CRE; Ai9-RCL^tdTomato^ (Vgat-CRE; Ai9)* double mutant mice and quantified, by immunostaining, the degree of colocalization between GABAergic neurons (tdTomato^+^) and MSK1^+^ cells in the cortex and striatum at postnatal ages P5, P10, P15 and P30 (Fig. 1C to 1H and Fig EV1 and EV2). The low percentage of tdTomato^+^ interneurons that are MSK1^+^ at P5 in the somatosensory cortex (≈32%) decreases sharply at P10 (≈21%), and until P30 (≈3%) (Fig. 1C and D and Fig. EV1). Around 85% of the MSK1^+^ neurons in the somatosensory cortex at P5 are GABAergic, a number that decreases to less than 40% at P30 (Fig. 1E).

In the striatum, there are ≈60% of MSK1^+^ cells at P5 and P10, and a reduction to ≈40% at P30 (Fig. 1G and Fig. EV2). Of the MSK1^+^ cells, approximately 96% are GABAergic neurons from P5 to P30 (Fig. 1F and H). Altogether, these results reveal that MSK1 is mostly restricted to GABAergic neurons in the somatosensory cortex and the striatum.

### Morphometric and densitometric analysis of striatal MSK1^+^ cells along postnatal development

To analyse changes in MSK1 immunoreactivity in various stages of the mouse brain development, we used MATLAB maps coding area (Fig. 2 and Fig. EV3) to determine the optical density (OD) values of MSK1^+^ cells, correlated with its topographical distribution along the striatum, as previously reported (37). Between P5 to P10 the number of MSK1^+^ cells per 1000μm^2^ decreases, although at P10 cells show significantly higher OD. At P15, the number of immunoreactive cells increases again to values higher than P5 and P10, while OD decreases to the same level as in P5. Finally, at P30 we found the highest amount of MSK1^+^ cells per 1000μm^2^ (Fig. EV3). OD values at this stage are lower than at P10, but higher than at P5 and P15 (Fig. EV3). The mean area of the immunoreactivity in MSK1^+^ cells keeps getting progressively bigger from P5 until P30 (Fig. EV3). We also observed changes in the distribution of OD values depending on age (Fig. 2). At P5, MSK1^+^ cells concentrate in low OD values, while at P10 they move to medium values, with few cells in either very low or very high OD. At P15 they go back to mainly low OD values, and at P30 the distribution gets more homogeneous, with cells in low, medium, and high OD (Fig. 2C). Together, these data show that *Msk1* expression is temporarily regulated during striatal development at two different stages, between P5 to P10 and between P15 to P30.

**Figure 2.**
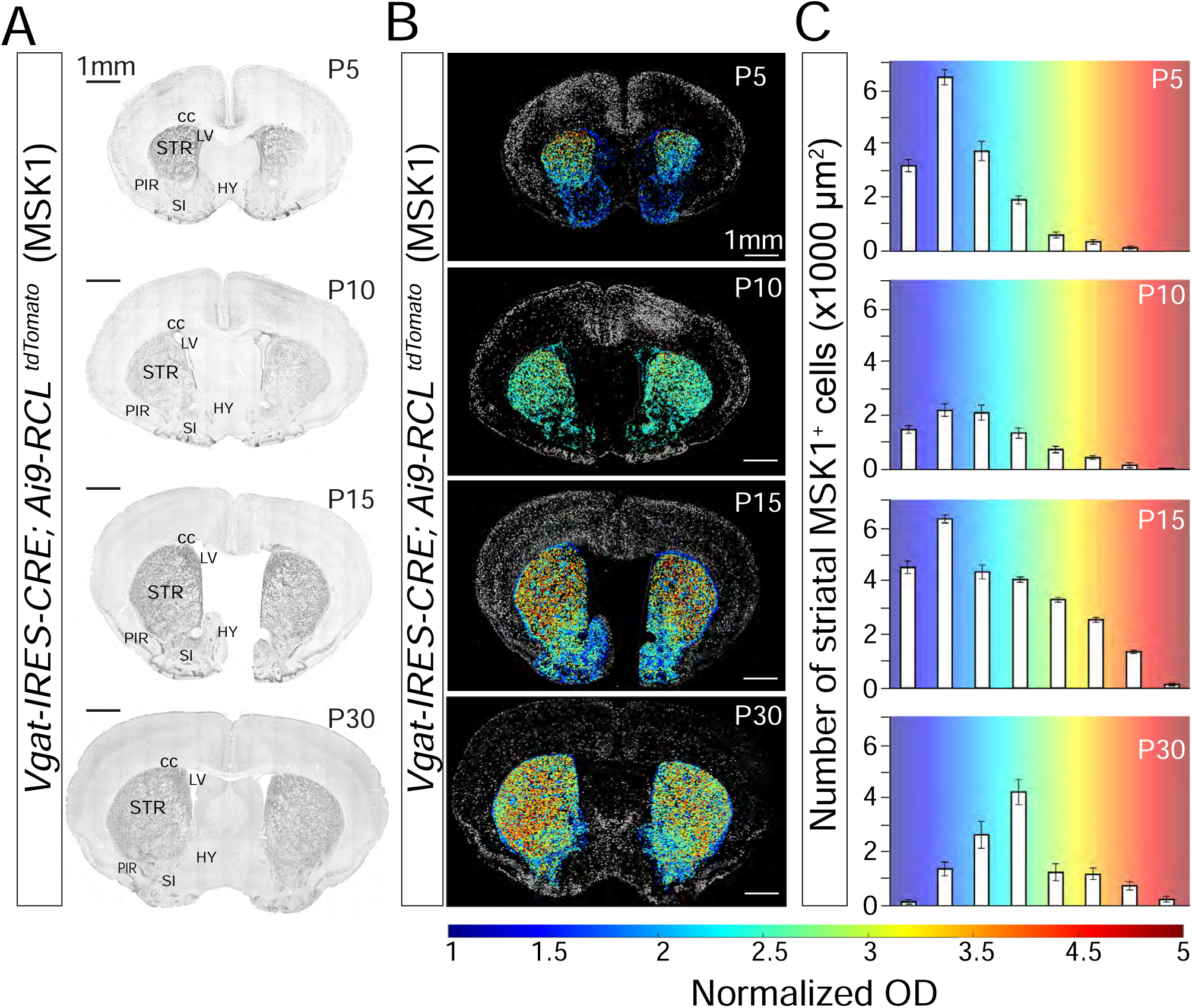
Developmental changes in MSK1^+^ cells in the mouse striatum. (A) IHC/DAB images showing MSK1^+^ cells in the brain at different stages of development (P5, P10, P15 and P30). STR=striatum, LV=lateral ventricle, HY=hypothalamus, SI=substantia innominata, PIR=piriform cortex, cc=corpus callosum. (B) MATLAB maps showing the distribution and area of MSK1^+^ cells in the striatum at different postnatal ages (P5, P10, P15 and P30). Cells from the striatum have been highlighted and show variations in OD. (C) Distribution of OD values (blue – low OD, red – high OD) from striatal MSK1^+^ cells.

### Generation and characterization of *Msk1 Exon IV* KO mice

To understand the role of MSK1 during mouse brain development and adulthood, we generated a new *Msk1* knock-out model. We removed the *Msk1* Exon IV with CRISPR/Cas9 to generate a premature STOP codon (TGA) in the Exon V (Fig. 3A and B). We targeted Exon IV, and not Exon I, due to the presence of an alternative ATG start codon in the Exon III of the gene that it is frame with the coding ORF. This ATG could, putatively, be used as an alternative to the one located in Exon I. We confirmed the edition by Sanger sequencing (Fig. 3A) and genotyped the offspring by two PCRs (Fig. 3C). The *Msk1 Exon IV* KO*– mice* (*Msk1^IV^* KO) are viable and fertile, with non-evident developmental problems or abnormalities at P30 such as different size or weight compared to control littermates, gross body tremors, hindlimb or forelimb clasping. We did not detect MSK1 protein levels by western blot (Fig. 3D) and immunofluorescence assays when using specific antibodies directed against the C-terminal domain (Fig. EV4A). Also, we found that *Msk1* transcript levels in these mice are significantly reduced compared to control mice (Fig. 3E), probably due to a nonsense-mediated RNA decay process derived from the gene editing (38).

**Figure 3.**
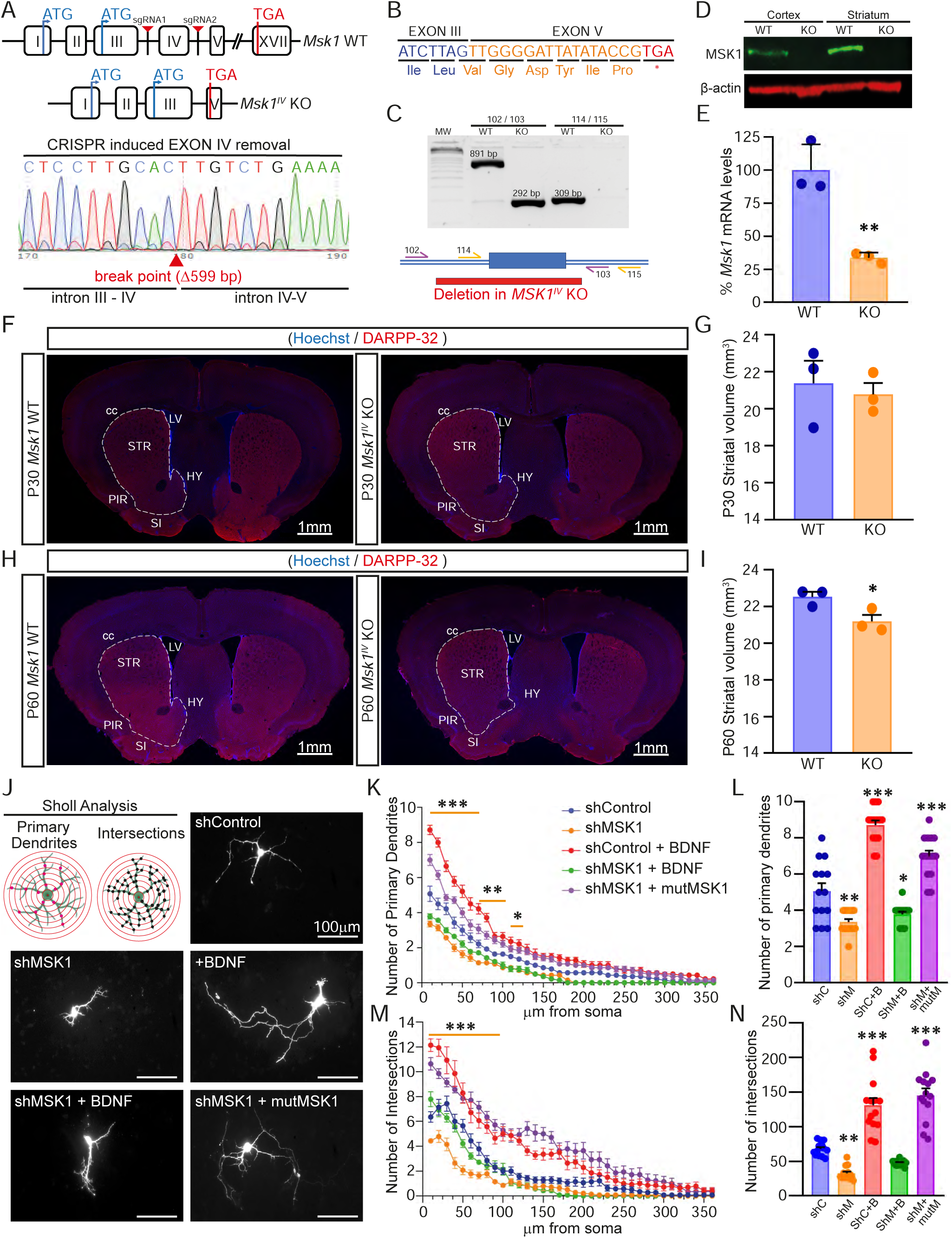
MSK1 controls postnatal striatal growth and arborization of striatal neurons. (A) *Msk1* Exon IV KO allele removal induced by CRISPR/Cas9 (top), and genomic Sanger sequencing (bottom) of the CRISPR/Cas9 edition result. (B) Exon IV removal creates a premature stop codon in Exon V, generating *Msk1^IV^* KO mice. (C) PCR and scheme of the genotyping of *Msk1^IV^* KO offspring using 2 different sets of primers (102/103 and 114/115). (D) Western Blot from cortical and striatal lysates of *Msk1^IV^* KO mice using specific monoclonal antibodies against the C-terminal domain of the protein. (E) Quantitative PCR of *Msk1* transcripts from mRNA extracted from the striatum of P30 *Msk1^IV^* KO mice. *\*\*P<0.01* (mean ± SEM; n=3; two tailed unpaired Student’s t test). (F and H) Representative coronal sections of wild-type and *Msk1^IV^* KO mice immunostained for DARPP-32 at postnatal ages P30 and P60. Dashes lines indicate the boundary of the striatum, determined with the Allen Brain Atlas, considering both stainings and the anatomical boundaries of the striatum. STR=striatum, LV=lateral ventricle, HY=hypothalamus, SI=substantia innominata, PIR=piriform cortex, cc=corpus callosum. Scale bars, 1mm. (G and I) Striatal volume estimation of *Msk1^IV^* KO compared to wild-type mice at P30 and P60 applying Cavalierís principle. \**P*<0.05 (mean ± SEM; n=3 mice, 10 to 12 sections per brain; two tailed unpaired Student’s t test). (J) Representative images of 7 DIV cultured wild-type striatal neurons 5 days after transfection with a reporter plasmid for EGFP containing a control shRNA (shControl), shRNAs against MSK1 (shMSK1) or shRNAs against MSK1 plus a resistant form of MSK1 against them (shMSK1+mutMSK1). BDNF (50 ng/ml) was added 2 days after transfection. (K to N) EGFP positive neurons were analysed by fluorescence microscopy and neuronal arborization was quantified using Sholl analysis. \**P*<0.05, \*\**P*<0.01, \*\*\**P*<0.001. (mean ± SEM; n=14 neurons per condition. Two-way ANOVA followed by Tukeýs multiple comparisons test, comparing each case against the control (shControl)).

### MSK1 mediates postnatal striatal growth and MSNs arborization

Using Cavalierís method, we found a significant difference in the volume of the striatum between wild-type and *Msk1^IV^* KO littermate mice at P60 (Fig 3F – 3I and Fig. EV4). However, there is not a significant reduction in the total length and weight between the brains of the *Msk1^IV^*KO and WT mice at this age (Fig. EV4B). Since the arborization of cultured striatal neurons relies on the BDNF-dependent MAPK pathway activation (20), we wondered whether MSK1 may mediate the growth effects of BDNF in MSNs during mouse brain development in a cell-autonomous way. To test this hypothesis, we transfected wild-type striatal cultured neurons with control or specific *Msk1* miRNAs-based shRNAs (shMSK1) (Fig. EV5). We found that the length, number of primary dendrites, and arbor complexity was reduced in the absence of MSK1, even under the presence of BDNF (Fig. 3J). Moreover, the total number of primary dendrites, neurite complexity and length were rescued by overexpression of a mutant form of *Msk1* (mutMSK1) resistant to the shMSK1 (Fig. 3J). These results show that MSK1 controls the BDNF-dependent arborization of mouse striatal MSNs and is required for normal striatal maturation.

### MSK1 mediates BDNF-dependent MeCP2 S421 phosphorylation

Stimulation of cortical and hippocampal neurons with BDNF induces phosphorylation of MeCP2 through the CaMKII pathway but independently of the MAPK pathway (25). MeCP2 is expressed by both glutamatergic and GABAergic neurons, and its activity is critical for proper brain development and striatal function (39). We hypothesized that MSK1 could also mediate the BDNF-dependent MeCP2 phosphorylation in striatal neurons. BDNF treatment led to an increase in pTrkB and pERK1/2 levels in cultured striatal neurons, but also to an increase in pMeCP2(S421) levels (Fig. 4A). The latter was reduced after inhibiting the MAPK pathway with U0126 (Fig. 4A) and also by the absence of MSK1 (Fig. 4B). Next, we tested if MSK1 and MeCP2 interact in the cell nucleus. Western blotting of HA immunoprecipitates, obtained from purified nuclear lysates of HEK293-FT cells expressing HA-MeCP2 and MSK1-myc, shows that both proteins interact independently of their phosphorylation state (Fig. 4C). Together, these results suggest that MSK1 forms part of a nuclear complex with MeCP2, and that in striatal neurons MSK1 is required for the BDNF-dependent MeCP2 S421 phosphorylation.

**Figure 4.**
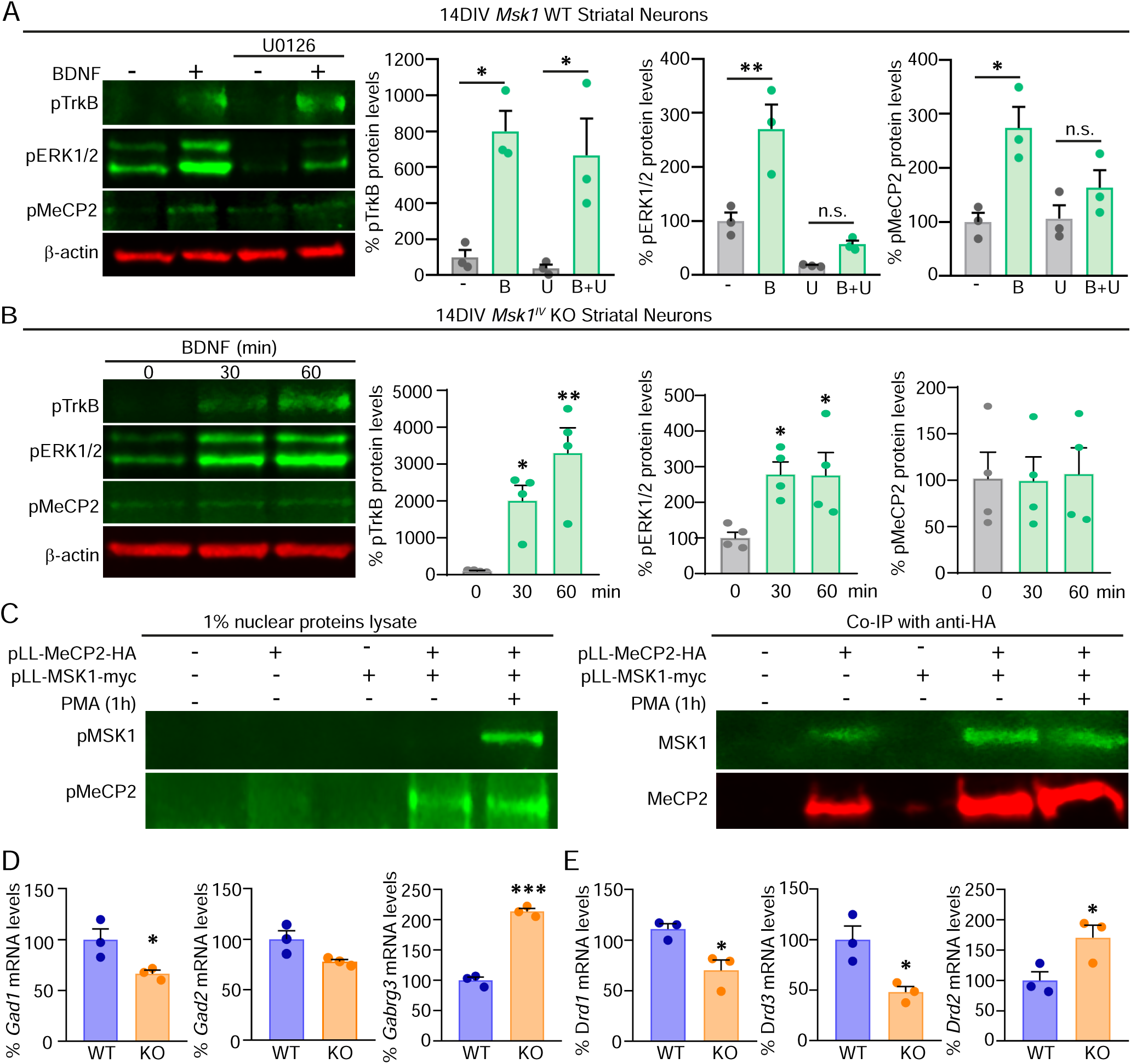
MSK1 interacts and mediates BDNF-dependent MeCP2 S421 phosphorylation. (A and B) Quantitative Western-Blot analysis of WT and *Msk1^IV^* KO cultured striatal neurons. Graphs values represent pTrkB, pERK1/2 and pMeCP2-pS421 levels relative to actin levels. (A) WT striatal cultured neurons with different treatments=1h BDNF (50 ng/mL) stimulation. U=30 min U0126 treatment (100 nM). B+U= 30 min pre-treatment with U0126 (100nM) followed by 1h BDNF stimulation (50ng/mL) n=3. (B) Striatal *Msk1^IV^* KO were treated with BDNF for 30- and 60-min. n=4. (C) Lysates from HEK293-FT cells co-transfected with expression plasmids for MeCP2-HA and MSK1-myc were immunoprecipitated with anti-HA antibodies coupled to magnetic beads. Western blots were performed to detect the interaction of MSK1 with MeCP2 before or after inducing MSK1 activation by PMA (200 nM; 1h). (D) qPCR analysis of mRNAs from P60 WT and *Msk1^IV^* KO striata for *Gad1*, *Drd1, Drd2, Drd3* and *Gabrg3*. n.s.= non-significant, \**P*<0.05, \*\**P*<0.01, \*\*\**P*<0.001 (mean ± SEM; two tailed unpaired Student’s t test).

### MSK1 regulates the expression of genes related to the GABA and dopamine system

The lack of *Mecp2* in the striatum alters gene transcription (39, 40). To address whether the transcriptional effects of MeCP2 were dependent on MSK1 we decided to study, among others, the mRNA levels of genes which expression is altered by the lack of MeCP2 in the striatum. We found by qPCR of cDNAs obtained from the striatum of P60 *Msk1^IV^* KO mice, a reduction in the expression levels of the gene glutamate decarboxylase 1 (*Gad1*), which codes for GAD67, and an increment in the expression of the specific MSNs GABA_A_ receptor subunit gamma 3 (*Gabrg3*), compared to those of wild-type mice (Fig. 4D). Moreover, while the mRNA expression of the dopamine receptors *Drd1* and *Drd3* was reduced, the levels for *Drd2* were significantly increased (Fig. 4E) (39, 41). Furthermore, we observed no changes in the expression of *Grin1*, (glutamate ionotropic receptor NMDA type subunit 1), *Psd95* (glutamatergic post-synaptic density protein 95) and the serotonin receptor *5-Htr2c* (5-Hydroxytryptamine Receptor 2C) (Fig. EV6). The lack of MSK1 in our mutant mice also affects the expression of the genes *Irak1* (interleukin-1 receptor-associated kinase 1)*, Satb1* (special AT-rich sequence binding protein 1)*, Dlk1* (delta like non-canonical Notch ligand 1) *and Pvalb* (parvalbumin) (Fig. EV6), as for mice in which *Mecp2* is removed only in GABAergic neurons (39, 40). We found no changes in neither the expression of the specific MSNs gene *Darpp-32* nor in the astrocytic gene *Gfap* (glial fibrillary acidic protein), but a reduction in the levels of *Ppargc1a* (peroxisome proliferator-activated receptor gamma coactivator 1-alpha) (Fig. EV6), whose expression is known to be dependent on MSK1 (36). Altogether, our data indicate that MSK1 is required for the expression of genes implicated on the functionality of the GABA and dopamine networks in the striatum, in a similar way to what has been described for MeCP2-deficient mice.

### Lack of MSK1 alters emotional behaviour

To investigate the behavioural consequences, if any, in our KO mouse model, we first decided to test the locomotor activity of males and females by the open field and the accelerating rotarod assays (Fig. 5A and B). Neither in the open field nor in the rotarod test they show obvious locomotion problems. We also evaluated the innate and specific social responses together with anxious and depressive behaviours on both sexes. We assessed innate responses by the nest building test, while used the three-chambered social test to analyse the specific social behaviour. When compared to control mice, both *Msk1^IV^*KO males and females showed poorer nest building at 4h and 24h (Fig. 5C). In addition, mutant mice of both sexes showed hypersocial behaviours, as they spent longer time sniffing for a conspecific mouse than control wild-type mice in the three-chambered social test (Fig. 5D). When analysing the anxiety response, we observed that *Msk1^IV^* KO males, but not mutant females, entered more and spent longer time in the centre of the open field (Fig. 5A). Also, by the marble-burying test, *Msk1^IV^* KO males, but not females, buried less marbles than controls (Fig. 5E). Finally, we observed that *Msk1^IV^*KO mice remained immobile for longer time than control littermates in the forced swimming test (FST), showing a depressive-like or demotivation behaviour (Fig. 5F). All together, these data indicate that the lack of MSK1 from embryogenesis alters adult mouse behaviour.

**Figure 5.**
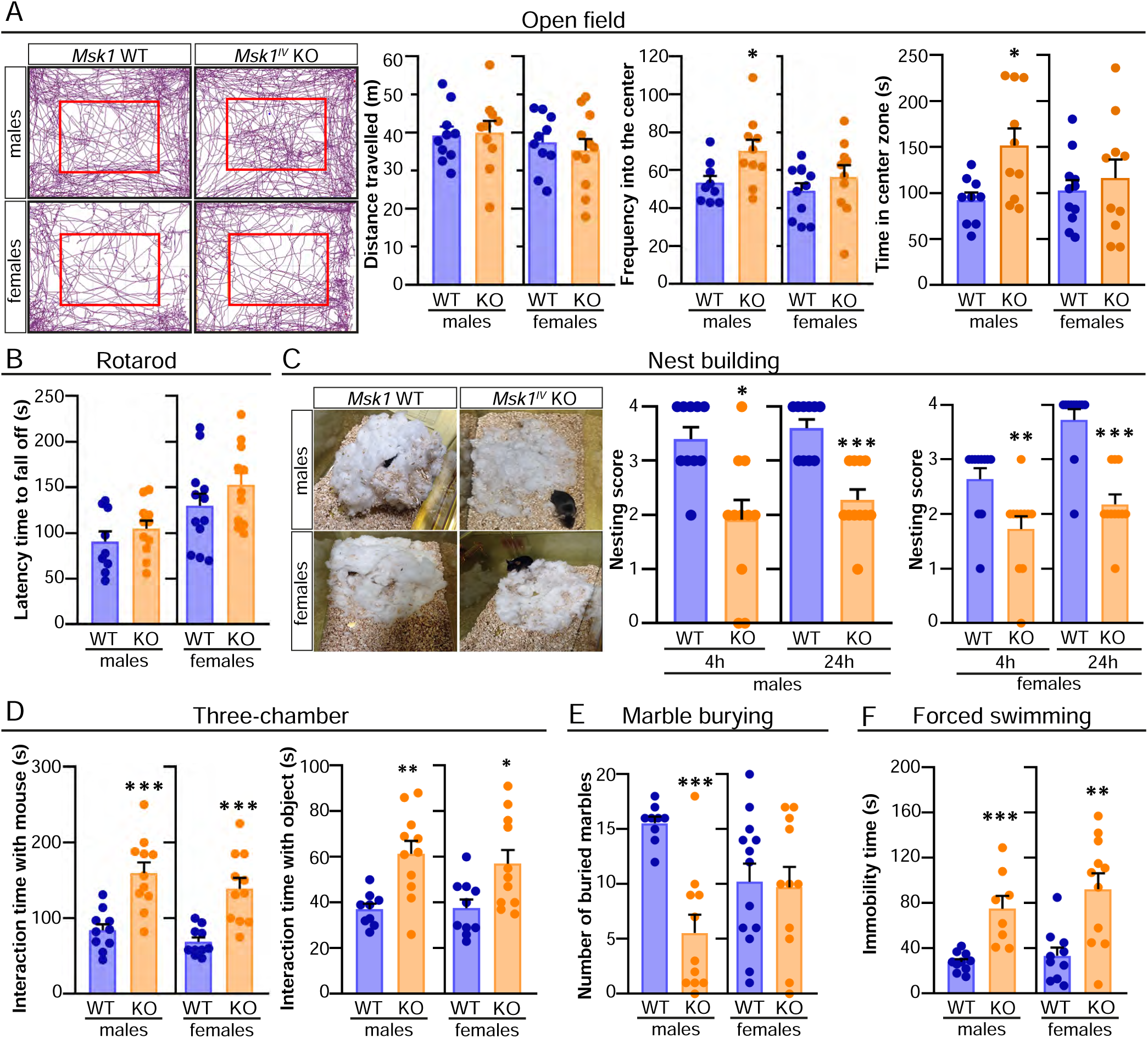
MSK1 deletion causes altered emotional behaviour. (A and B) Locomotor activity of adult *Msk1^IV^* KO mice compared to control wild-type mice. (A) Distanced travelled, frequency into the center and time spent in the center zone measured in the open field. (B) Latency time for the mice to fall off in the rotarod test. (C) Representative pictures of a nest built after 24h and nesting score in the nest building test at 4h and 24h. (D) Interaction time with conspecific mouse and with object in the three-chamber test. (E) Anxiety response assessed by the marble burying test result after 30 min performance. (F) Depressive-like state was analysed by the immobility time of mice during the Forced swimming test. In all cases male and female WT and *Msk1^IV^* KO animals were 2 to 4 months old and statistical analysis was performed with two tailed unpaired Student’s t test \**P*<0.05, \*\**P*<0.01, \*\*\**P*<0.001. (mean ± SEM; n= 9 to 13 mice per condition and sex).

## Discussion

Our study uncovers a new molecular mechanism through which the lack of MSK1, which expression we found to be controlled during postnatal development in GABAergic neurons, impacts the arborization of MSNs and the growth of the striatum. This absence leads to the lack of phosphorylation of MeCP2 and alters the expression of genes related to GABAergic and dopamine responses in the adult mouse striatum. These abnormalities are linked to atypical innate and social responses, as well as anxiety and depression-like behaviours in mice. Our findings support the notion that dysfunction in the striatum, due to developmental abnormalities caused by the lack of MSK1, might play a central role in the pathophysiology of schizophrenia.

### Msk1 expression is controlled during development

It has been well studied that MSK1 is mainly located in striatal MSNs and Purkinje neurons in the cerebellum of the adult mouse brain (33–36, 42), but much lower in the brain cortex and hippocampus (33, 35). Our results demonstrate that MSK1 in the early postnatal somatosensory cortex and striatum is mostly restricted to GABAergic neurons. Also, we observed that its expression is differentially regulated in both areas during postnatal development. While *Msk1* expression decreases in the somatosensory cortex, it increases in striatal cells until young adulthood. This suggests that there may be distinct responses of GABAergic neurons in both the striatum and cortex to stimuli that controls MSK1-dependent neuronal regulation events. Furthermore, the role of this kinase may vary depending not only on the developmental stage but also on the specific brain region and neuronal population in which it is expressed. However, it remains unclear whether MSK1 activity can be controlled by the same stimuli in both neuronal populations.

*Msk1* is primarily expressed by MSNs in the adult striatum (36), so these should be the most affected by the lack of MSK1 during postnatal brain development. Of note, our morphometric and densitometric results indicate that MSK1 levels are not equal, neither constant, in all striatal cells during postnatal development. While this may indicate a different maturation stage of the major MSNs populations, that express *Drd1* or *Drd2* respectively (43), it is also possible that the maturation of the dorsal and ventral striatum differs due to the regulation of MSK1-dependent genes. More detailed experiments should be done in the future to test this hypothesis.

#### MSK1 role in postnatal maturation of striatum and MSNs arborization

Many studies have described that BDNF controls axonal and dendritic arborization of neurons depending on their class (17, 18, 23, 44–47). In terms of neuronal arborization, some reports have shown that pyramidal neurons located in layer IV are more responsive to BDNF that those located in layer VI (44, 45). Other authors have shown that BDNF promotes arborization growth in GABAergic neurons but not in glutamatergic neurons in the hippocampus (17). Studies with mice lacking BDNF in the CNS revealed an area-specific requirement for dendritic growth of striatal neurons but not for the growth of CA1 pyramidal neurons, suggesting that the differential responsiveness to BDNF depends on neuron-specific programs (18, 19).

The BDNF/MAPK pathway is required for the proper growth of MSNs as well as for the postnatal growth of the striatum (18, 20, 48). BDNF mediates the activation of the RSK (RSK1, RSK2, RSK3 and RSK) and MSK (MSK1, MSK2) family members (49–51). Our data on *Msk1* developmental expression in the striatum raise the possibility that it controls the specific growth of MSNs. Indeed, we observed that the lack of MSK1 clearly affects the postnatal growth of the striatum. Moreover, we found that the specific downregulation of MSK1 in striatal cultured neurons leads to a significant poor arborization even in the presence of BDNF. RSKs and MSKs can be activated by similar signals, suggesting some redundancy in the phosphorylation of substrates *in vitro* (51, 52). However, RSKs and MSKs are localized in different cells localization in the adult mouse brain (33, 34). Our results reveal postnatal regulated expression of *Msk1* in GABAergic interneurons and MSNs, suggesting that the functions of the RSK and MSK family members could not be totally redundant despite of what has been previously indicated (51).

#### Altered gene expression and behavioural features in MSK1^IV^ knockout mice

Our results demonstrate that MSK1 interacts and controls BDNF-dependent MeCP2 S421 phosphorylation in striatal neurons. These results seem to differ from those others that show that MAPK signalling does not mediate MeCP2 phosphorylation, but it is dependent on the activation of CaMKII by BDNF (25). Of note, *Mecp2* is expressed not only by glutamatergic but also by GABAergic neurons in different brain areas (13). In addition, MeCP2 regulates the expression of the genes coding for the GABA synthesizing enzymes GAD65 and GAD67 (53, 54). Therefore, we expect that the discrepancy between our results and the ones mentioned, could be due to the different type of neurons that we are analysing, striatal neurons, instead of the ones analysed in those studies, cortical and hippocampal neurons.

We found that the absence of MSK1 alters striatal expression of *Gad1*, *Drd1*, *Drd2*, *Drd3*, *Pvalb* and *Irak1*, similarly to what has been observed in mice lacking MeCP2 in GABAergic neurons (40, 53, 54). These data show that MSK1 influences the GABA and dopamine system in the striatum. However, the lack of alterations on the expression of *Psd95* and the serotonin receptor *5-Htrc-2* by the MSNs, indicate that MSK1 may not be involved in the regulation of the glutamatergic afferent projections that these neurons receive from cortical areas, neither in the serotonin-dependent functions. MSK1 regulates gene expression by modulating the phosphorylation of Histone H3 (55), transcription factors such as CREB (28, 56), and DNA binding proteins (57, 58). Our experimental evidence suggests that MSK1-dependent MeCP2 S421 phosphorylation also modulates genes involved in GABA and dopamine functions in the striatum. Despite its role as regulator of MeCP2 activity, MSK1 depleted mice do not show Rett syndrome-like features but resemble some of those observed in mice lacking MeCP2 in MSNs. At their core, behavioural dysfunctions observed in MeCP2 null mice will most likely be due to a combination of alterations of glutamatergic and GABAergic circuits in different brain areas including striatum.

*Msk1^IV^* KO mice display some features linked to striatal and dopamine functions that are also observed in schizophrenia patients (59). There are several mouse models in which it has been shown that dysfunctions in the development of the GABAergic network (60) and the dopaminergic system (61) are implicated in SCZ. The reduction in *Drd1* and increment on *Drd2* mRNA levels in our *Msk1^IV^* KO mice, indicates a possible disruption on the dopamine signalling. MSNs constitute two different pathways that controls movement: the direct pathway, controlled by DRD1 expressing MSNs (D1-MSNs), implicated in movement promotion, and the indirect pathway, controlled by DRD2-MSNs (D2-MSNs) inhibiting movement. Surprisingly, our mice do not show motor impairments, despite the disruption in the mRNA levels of both dopamine receptors and the striatal growth. In a transgenic mouse model, it has been observed that overexpression of *Drd2* results in hyperactivity, primarily caused by a reduced rate of dopamine reuptake in the synaptic space (62). It is important to note that the levels of DRD1 remain unaffected in these mice in spite of *Drd2* overexpression due to the transgene. It is possible that the imbalance in dopamine receptors expression observed in *Msk1^IV^* KO mice may not be enough to cause alterations on the motor behaviour. Further studies on the activity and regulation of dopamine receptors should be done to investigate the mechanisms that enable normal movement in MSK1-deficient mice, despite the different expression of dopamine receptors in the striatum.

Similar to *Erbb4-*deleted and *Glud1*-deficient mice (60, 63), which are schizophrenia-like mouse models, nesting behaviour is disrupted in *Msk1^IV^* KO mice. This suggests that our mice exhibit self-neglect attitudes, which are characteristic negative symptoms observed in schizophrenia patients (64). Other negative symptoms, such as depression, demotivation, and avolition, are typically assessed using the FST in mice (65). In this test, *Msk1^IV^* KO mice display increased freezing time, indicating depressive and demotivated behaviour.

Additionally, *Msk1^IV^* KO male mice bury significantly less marbles compared to control littermates, like *Erbb4* mutant mice (60). This result is consistent with the number of entries in the centre zone observed in the same group in the open field, as both results resemble for less anxiety.

In addition, *Msk1^IV^* KO mice are overly social compared to controls, as they spend more time interacting with conspecific. This is a characteristic of some neuropsychologic illness mouse models like *Grin1* KO, a schizophrenia-like mouse model (66). Hypersociability is related with a reduction in the source of BDNF from the Ventral Tegmental Area (VTA) to the Nucleus Accumbens (NAc), part of the ventral striatum involved in mediating motivational and emotional processes (67). The lack of MSK1 in the MSNs of the NAc could alter them similarly to the lack of BDNF signalling arriving from the VTA. Due to this, MSK1 absence would lead to hypersocial behaviour, as observed in the *Msk1^IV^* KO mice.

Our study reveals different behaviours between male and female *Msk1^IV^*KO mice, reflected in both the frequency in entering to the centre of the open field, as in the number of buried marbles. Sexual hormones are known to have an important role in brain development and disruptions in them are linked to neurological disorders such as SCZ. For example, low circulating estradiol levels are associated with female schizophrenia patients (68, 69). In addition, altered GABA signalling is related to SCZ as it has been shown that there is a lower expression of GABA-related genes in some brain areas of male schizophrenia patients in contrast with higher levels in the same areas of female patients (70). All of this, together with the earlier onset of the disorder in males than in females (71) and the possible role for MSK1 in brain development and GABA signalling, could explain some of the differences in our results.

The effects of sex on schizophrenia are multifaceted and complex, encompassing variations in symptom manifestation, social outcomes, and responses to treatment between men and women. Understanding these differences is of utmost importance in order to develop more effective and personalized treatments for individuals living with schizophrenia.

Our data highlights the view that MSK1 modulation may open new therapeutic approaches and provide mechanistic insights into the cell autonomous regulation of GABAergic development through gene expression control, and the potential link of MSK1 with schizophrenia-like behaviours. We remark the importance of considering the differences between sex in order to address future research.

## Methods

### Animals

All of the experiments using mice were conducted with permission from the Consejería de Agricultura, Ganadería y Desarrollo Rural of Castilla y León Government, after approval by the ethical committee of the University of Salamanca and following the European Directive 2010/63/EU and the Spanish Royal Decree 53/2013-Law 32/2007. All animals were kept on a 12:12 light/dark cycle, receiving food and water *ad libitum*. *Vgat-CRE; Ai9* mice were generated by breeding the mouse lines Vgat-IRES-CRE (B6J.129S6(FVB)-*Slc32a1^tm2(cre)Lowl^*^/MwarJ^; JAX stock #028862; Vong et al., 2011) with mice Ai9 (B6.Cg-*Gt(ROSA)^26Sortm9(CAG-tdTomato)Hze^*^/J^; JAX stock #007914; Madisen L et al., 2010). Founder (F_0_) *Msk1^IV^* KO mouse was bred with C57BL/6 mice obtained by local breeding, and heterozygous offspring were back-crossed to C57BL/6J mice for 4 generations before establishing the mouse line *Msk1^IV^*KO (B6-Rps6ka5^em3SAL^).

### Quantitative Real-Time PCR (qPCR)

cDNA was synthesized using the RevertAid retrotranscriptase (ThermoFisher; Manufacturer instructions) from 0.5μg of total RNA isolated from cortex or striatum using Trizol (Invitrogen; Manufacturer instructions). qPCR was performed using Sybr^TM^ Green PowerUp^TM^ mix (Applied Biosystems^TM^; Manufacturer instructions) and specific primers for *Msk1, Gad1, Gad2, Gabrg3, Drd1, Drd2, Drd3, Grin1, Psd95, 5-Htrc-2, Irak1, Satb1, Dlk1, Pvalb, Darpp-32, Gfap, Ppargc1a* and *Tbp* (see Table 1) on a QuantStudio^TM^ 7 Real-Time PCR System (ThermoFisher). Data were analysed using the 2^-ΔΔCt^ method.

**Table 1:**
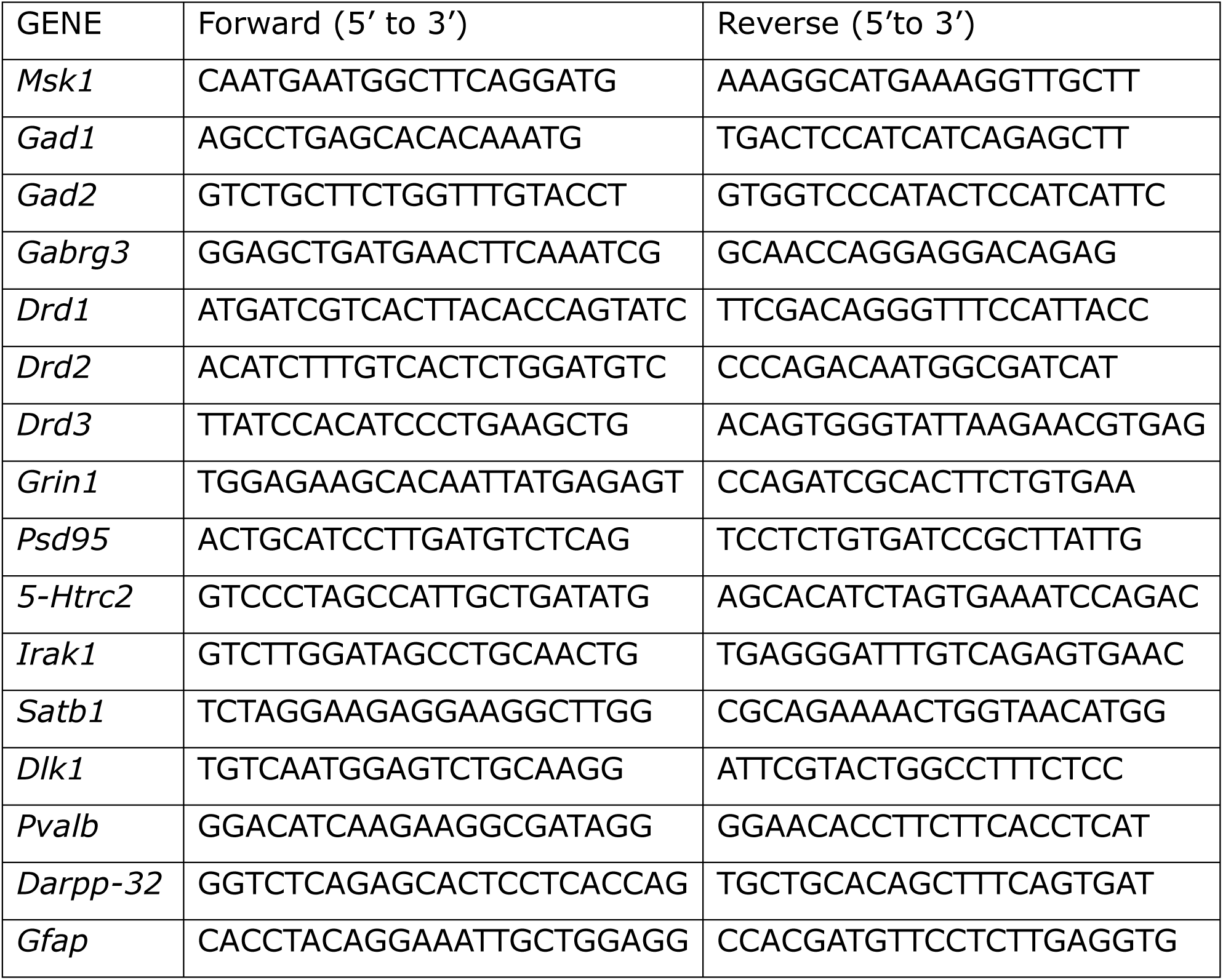
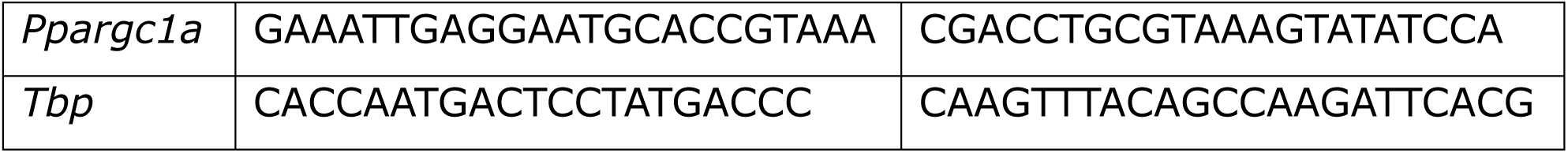
Primers used for qPCR assays.

### Quantitative Western Blot

Total protein lysates were obtained in RIPA buffer (Tris-HCl 25mM pH7.5, NaCl 150mM, TritonX-100 1%, sodium deoxycholate 1%, and SDS 0.1%) containing protease and phosphatase inhibitors (Pierce, Thermofisher) and quantified by the BCA method (Pierce Protein Assay, Thermofisher; Manufacturer instructions). Equal amounts of total protein (25-35μg) were loaded and separated by electrophoresis in Bis-Tris 10% acrylamide gels. After, proteins were transferred to Immobilon-FL PVDF membranes (Merck-Millipore) and washed twice in TBS buffer followed by incubation in Intercept-TBS Blocking Buffer (Li-COR; Manufacturer instructions). Antibodies and concentrations were as follow: rabbit monoclonal anti-MSK1 (C27B2; Cell Signaling) 1:1000, mouse monoclonal anti-β-actin (669-1-Ig; Proteintech) 1:20000, mouse monoclonal anti-phospho-pan-Trk (#9141; Cell Signaling) 1:1000, rabbit polyclonal anti-MeCP2 phospho S421 (ab254050; Abcam) 1:500, rabbit monoclonal anti-pERK1/2 (#9101; Cell Signaling) 1:3000, mouse monoclonal anti-Myc (sc-41; SantaCruz) 1:200, mouse monoclonal anti-MeCP2 (Quimigen 10861-1-AP) 1:500; rabbit polyclonal anti-MSK1 phosho Thr581 (#9595; Cell Signaling). Fluorescent secondary antibodies used were: IRDye 800CW Donkey anti-rabbit IgG (926-32213; Li-COR) 1:20000 and IRDye 680LT Donkey anti-mouse IgG (926-68022; Li-COR) 1:20000. Images were capture using a Li-Cor Oddysey XF system. Fluorescent images were acquired using an Odyssey XF Imaging system (Li-Cor) and densitometry analyzed by Empiria Studio^TM^ Software.

### Immunohistochemistry

Mice were deeply anesthetized with sodium pentobarbital (60mg/kg body weight; i.p.), and perfused transcardially, first with ice-cold PBS 1X (70mL) followed by the same volume of ice-cold PFA 4% in PBS. Brains were extracted and post-fixed for 3h in PFA 4% at 4°C before immersion in sucrose 30% at 4°C for 48h. Afterwards, 40µm-thick coronal brain sections were obtained using a cryotome (Thermo Scientific ™, Microm™ HM 430). For colocalization, densitometric, and volumetric assays every 10^th^ section was used. Brain slices were collected in antifreeze solution (Glycerol 30%, Ethylene Glycol 30%, in PBS) and stored at −20°C until use.

For immunofluorescent assays, brain slices were rinsed twice in PBS 1X followed by a 10 min permeabilization step with Triton X-100 0.25% in PBS, and a 2h blocking step in a solution containing Triton X-100 0.3%, normal donkey serum 10% (0.17-000-121: Jackson ImmunoResearch) and bovine serum albumin 5% (BSA, MB04602; NZYtech) in PBS. After, slides were incubated for 24 hours at 4°C with MSK1 and DARPP-32 antibodies diluted in antibody solution (Triton X-100 0.3%, normal donkey serum 5% and BSA 1% in PBS). After, the slices were washed 4 times in PBS 1X at RT for 15 min, followed by 1h incubation in the dark with antibody solution containing the secondary antibodies and 4μM Hoechst 33258. After 3x 10 min washes in PBS, tissue slices were mounted onto Superfrost glass-slides previously coated with gelatine solution (Tris-HCl 50mM pH7.5, gelatine from porcine skin 0.2 %) and let dry at RT protected from light. The sections were after glass covered using a Mowiol/DABCO mounting solution (Mowiol 4-88 0.1%, glycerol 0.26%, DABCO 2.5%, Tris-HCl 0.1M pH8.5). Finally, the slides were kept protected from light at 4°C until visualization.

For densitometry assays, brain slices were rinsed twice in Phosphate Buffer (PB) 0.1M followed by a 10 min incubation with H_2_O_2_ and Methanol (1 part of Methanol, 1 part of H_2_0_2_ 30% and 8 parts of PB 0.1M). Later, the slices were washed 3 times in PB 0,1M for 5 min each at RT, followed by other 2x 5 min washes in TBS-T solution (Triton X-100 0.3% in TBS). Then, the slices were incubated for 24-48 hours at 4°C with antibody solution containing MSK1 antibody. After, the slices were rinse twice in TBS-T solution and incubated for 2h at RT with anti-mouse biotinylated secondary antibodies (BA-2000; Vector) diluted 1:200 in antibody solution. Sections were washed 2x 5 min in TBS-T and incubated for 3h at RT in avidin/biotin-peroxidase solution (PK-4000, Vectastain), followed by 2x 5 min washes with Tris-HCl 0.05M before been placed for 25 min in developing solution (3,3-diaminobenzidine tetrahydrochloride, DAB; Sigma-Aldrich, D-9015) with 0.006% H_2_O_2_. Slides were washed twice with a Tris-HCl 0.05 M solution and mounted onto Superfrost glass slides previously coated with 0,2% gelatine and let dry overnight. Finally, sections were dehydrated 3x in ethanol 100% for 1 min and 2x in xylol for 3 min, glass covered using Entellan^TM^ and kept at RT until visualization.

The following primary antibodies were used: rabbit monoclonal anti-MSK1 (C27B2; Cell Signaling) 1:1500, rabbit monoclonal anti-DARPP-32 (271111, Santa Cruz) 1:500. The secondary fluorescent antibodies used were donkey-anti-rabbit-Alexa 488 (21206; ThermoFisher) 1:400, donkey-anti-mouse-Alexa-555 (31750; ThermoFisher) 1:400.

### Immunofluorescence quantification

We selected representative slices of murine somatosensory cortex and striatum from each postnatal age (n=3 animals/age-group). High-resolution microphotographs (1024×1024 pixel resolution and 400Hz acquisition speed) were taken at 20X magnification using an inverted Stellaris 8 confocal microscope (Leica Microsystems, Wetzlar, Germany). Same laser gain, intensity, pinhole, and detection filter settings were maintained. In order to count the number of MSK1 and tdTomato expressing neurons and cell co-localization from both channels, images were analyzed using ImageJ software. From each region of interest (upper layers of the somatosensory cortex and dorsoventral striatum, from both hemispheres) a total of 6-9 images were captured and semiautomatically thresholded in order to segment cells from the background, enabling their visualization, quantification, and plotting. Firstly, image threshold was established for each channel using the Max Entropy/IsoData filter, and binary masks were generated and refined. Particle detection based on size was performed in order to identify positive cells. Co-localization results were obtained using a Boolean operator that adds two binary masks (from two different fluorescent labels of the same image) and returns a new one containing only pixels present in both previous masks. Finally, the number of MSK1^+^ and tdTomato^+^ particles were determined and measured.

### Densitometry quantification

High-resolution microphotographs (digital resolution of 132pixel/100µm^2^) were taken at 10X magnification under a Leica DMRX microscope with an MBF camera (MBF Bioscience CX9000; Williston, VT, USA) to prepare whole-section digital mosaics using the Neurolucida software (NL-Vs 8.0, MicroBrightField®, Inc., Williston, VT, USA). To set homogeneous microscopic illumination conditions and to calibrate optical density (OD) measurements, photographs were taken using a standardized grayscale range and a stepped density filter (11 levels) (®EO Edmund industrial optics-ref 32599, Karlsruhe, Germany). For morphometrical analysis, immunoreactive neurons were segmented by density thresholding using ImageJ software and the Maximum Entropy algorithm. Coordinates, area, and O.D. values of the segmented cells were collected to perform statistical analysis and to build MSK1 expression maps (MATLAB). O.D. was used as measurement of the intensity of the immunoreactivity of each cell. Normalized O.D. values of immunoreactive cells were calculated by subtracting the mean O.D. of the whole section from the O.D. values of segmented particles, divided by the gray standard deviation of the whole slice. The number of segmented cells was normalized to N/10,000μm^2^ of surface area. Extensive full-sections maps encoding the location, area and density of segmented particles were made using MATLAB software (MATLAB R-2017, Scatterplot function). O.D. was converted into color with the MATLAB color scale Jet from the Scatterplot function.

### Msk1^IV^ KO mouse line generation

The *Msk1^IV^* KO mouse line was generated by CRISPR/Cas9 edition in a C57BL/6J background. We designed specific crRNAs using the *Breaking Cas* application from the Spanish National centre of Biotechnology (https://bioinfogp.cnb.csic.es/tools/breakingcas/). Two specific crRNAs (Table 2) flanking *Msk1* Exon IV were selected for its removal. Previous to the microinjection, 5μL of each crRNA (250ng/μL) and 10μL of tracrRNA (250ng/μL) were annealed together to form the sgRNAs. To form the ribonucleoprotein complexes, 1μL of each sgRNAs was thereafter mixed with 1.2μL (300ng/μL) of *S. pyogenes* Cas9 (Integrated DNA Technologies) in 9.2μL of microinjection buffer (Integrated DNA Technologies) and incubated for 15min at RT. The Cas9/sgRNAs complexes were microinjected using borosilicate needles into the biggest pronucleus of one-cell C57BL/6J mouse zygotes at the Transgenic Facility (NUCLEUS, platform for research support of the USAL/IBSAL). The injected zygotes were kept at 37°C and 5% CO_2_ in KSOM media until the next day, when 15 to 20 viable 2-cell embryos were implanted into pseudo-pregnant CD-1 female mice. The offspring was characterized by PCR (Table 3; MSK1 Forward and MSK1 Reverse primers) and the promising PCR products were analysed by Sanger sequencing in order to determine the modification generated by the system. Founder animals (F_0_) were back-crossed to C57BL/6J wild-type mice to generate F1 *Msk1^IV/+^* mice. To detect the absence or presence of the mutant allele we used the primers 102/103 and 114/115 (Table 3).

**Table 2:**
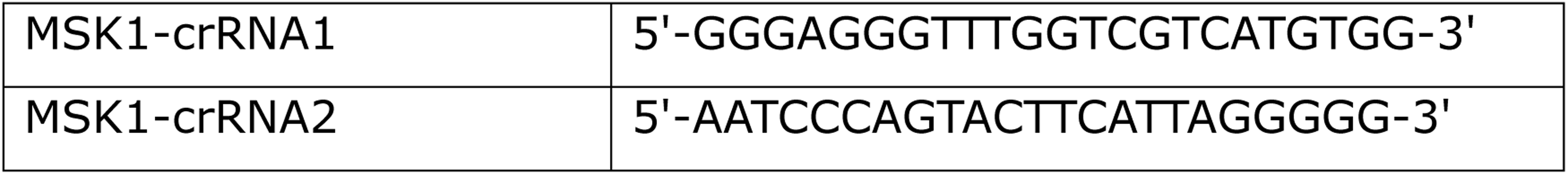
crRNAs used to generate the deletion of EXON IV.

**Table 3:**
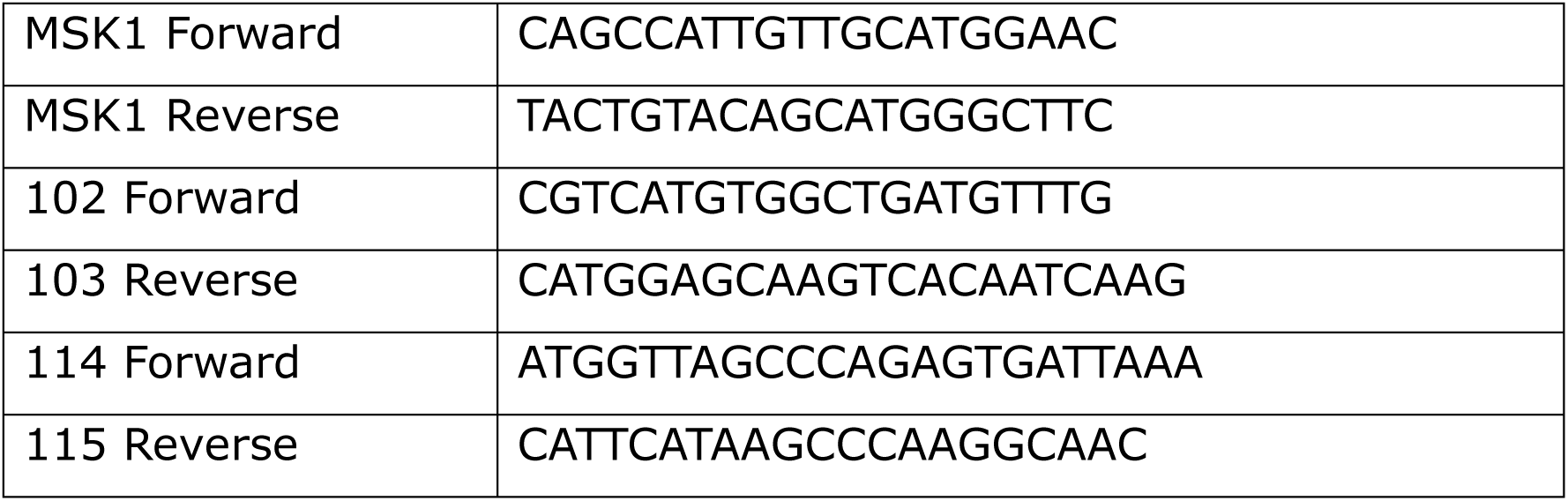
Primers for genotyping *Msk1^IV^* KO mice.

### Striatal volumetric analysis

Each brain was totally coronal sectioned using a cryotome (40μm thickness) and divided in 10 consecutive series. After identifying the correct series containing the first coronal cut of the striatum (Bregma +1.845mm; Allen Brain Reference Atlas) we did immunostaining assays as previously described using anti-DARPP-32 antibodies. Nuclei were counterstained with Hoechst 33258. Single plane tile scan confocal images were acquired at a 1024×1024 pixel resolution on an inverted Leica Stellaris 8 confocal microscope. 9 to 12 sections covering the striatum along the rostrocaudal axis were imaged using the same laser powers, gain, and detection filters. Both hemi-striata from each acquire image were delineated using the Allen Brain Reference Atlas as a guide and the area calculated using the ImageJ-Fiji software. To estimate the total volume, we applied the mathematical formula of Cavalieri’s principle 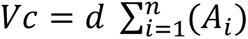, considering the distance between each brain slice (d), the calculated area of the striatum from each coronal section (A_i_) and the number of sections which contain the striatum (n).

### Primary striatal cultures

Striatal cultures were prepared from E14.5 – E16.5 mouse striata dissected in PBS-GB (D-Glucose 0.5%, BSA 0.1 %, Penicillin 10 μg/mL, Streptomycin 10 μg/mL). Tissue was dissociated by incubation with Trypsin-EDTA (0.25 %) solution (25200056; Gibco) at 37°C for 30 min, gently inverting the tube 3 to 4 times every 5 min, followed by mechanical disaggregation with a 10mL plastic pipette. Dissociated tissue was then diluted 1:1 with trypsin neutralizer solution (Neurobasal medium (Gibco), glutamine 0.2 mM (Gibco), NCS21 2% (Capricorn Scientific), trypsin inhibitor 0.035% (STEMCELL), DNase I (Sigma-Aldrich), resuspended with a 10 mL pipette, filtered through a 40 μm cell strained (BD) and centrifuge for 5min at 800g and RT. Cell pellet was thereafter resuspended in supplemented Neurobasal media (NCS21 2%, Penicillin 10 μg/mL, 10 μg/mL, glutamine 0.2 mM, HEPES 10 mM) and cells were plated at a density of 0.3×10^5^ cells/cm^2^ onto cell culture vessels or glass coverslips treated overnight with a solution containing Poly-D-Lysine hydrobromide (100 μg/mL; 70000-150000 MW; Sigma-Aldrich) and Laminin (10 μg/mL; CF858, Corning).

### HEK293-FT cell cultures

Cells were grown in supplemented DMEM (21699035; Gibco) (FCS 10%, glutamine 2 mM, HEPES 10 mM, Penicillin 10 μg/mL, Streptomycin 10 μg/mL, MEM non-essential amino acids 1X, 2-mercaptoethanol 0.001%). Cells were maintained at 37°C, 5 % CO_2_ and 95% humidity. Cultures were tested regularly for mycoplasma (6601; TaKaRa PCR Mycoplasma Detection Set).

### Plasmid generation

All vectors are based on the previously described dual-promoter lentiviral vector pLL-hSYN-DsRED-hSYN-EGFP (74), which basically contains two human Synapsin promoters able to transcribe efficiently and independent from each other the cDNAs located downstream of them. The Synapsin promoter is highly specific for neurons in primary cultures of cortical neurons.

#### Generation of shRNA expression vectors

The shRNA expressing vectors were generated as previously described (73). The specific miRNA-based shRNA against the mouse open reading frame (ORF) for MSK1 were generated using the BLOCK-iT™ RNAi Designer software (Invitrogen) and cloned into the pcDNA6.2-GW/miR vector according to instructions provided by the manufacturer. Positive clones were sequenced by PCR, and the miRNA-based shRNA cassettes, resulting from digestion with the restriction enzymes BamHI and NotI, were cloned into the pLL-hSYN-DsRED-hSYN-EGFP vector (74) or clone together into the pcDNA6.2-GW/miR vector into the BglII and NotI following manufacturer instructions. Positive clones including the 4 miRNA-based shRNAs were confirmed by Sanger sequencing before been clone into the pLL-hSYN-DsRED-hSYN-EGFP vector to generate the pLL-shMSK1 vector. The oligonucleotides used to generate the miRNA-based shRNAs are listed in Table 4.

**Table 4:**
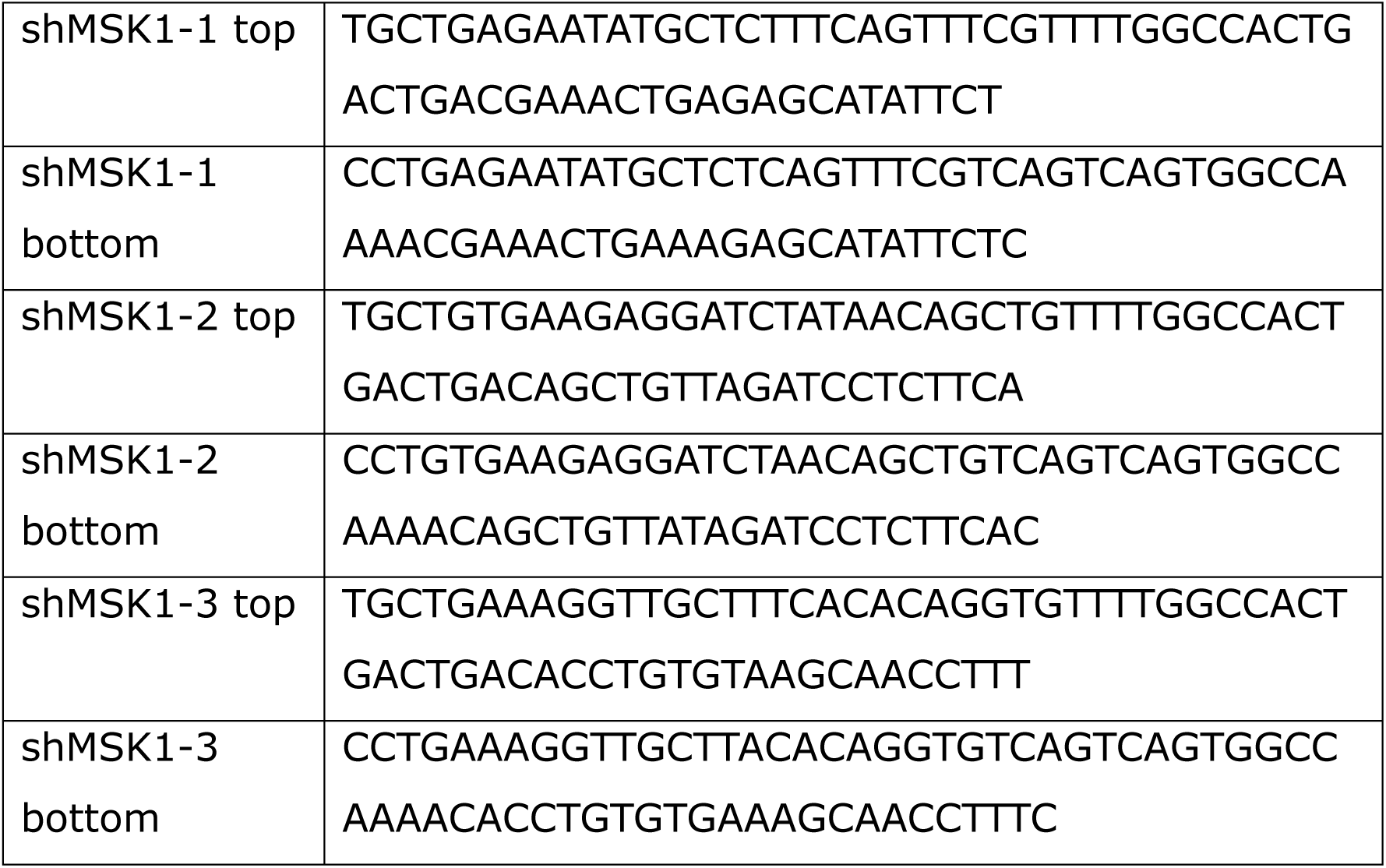
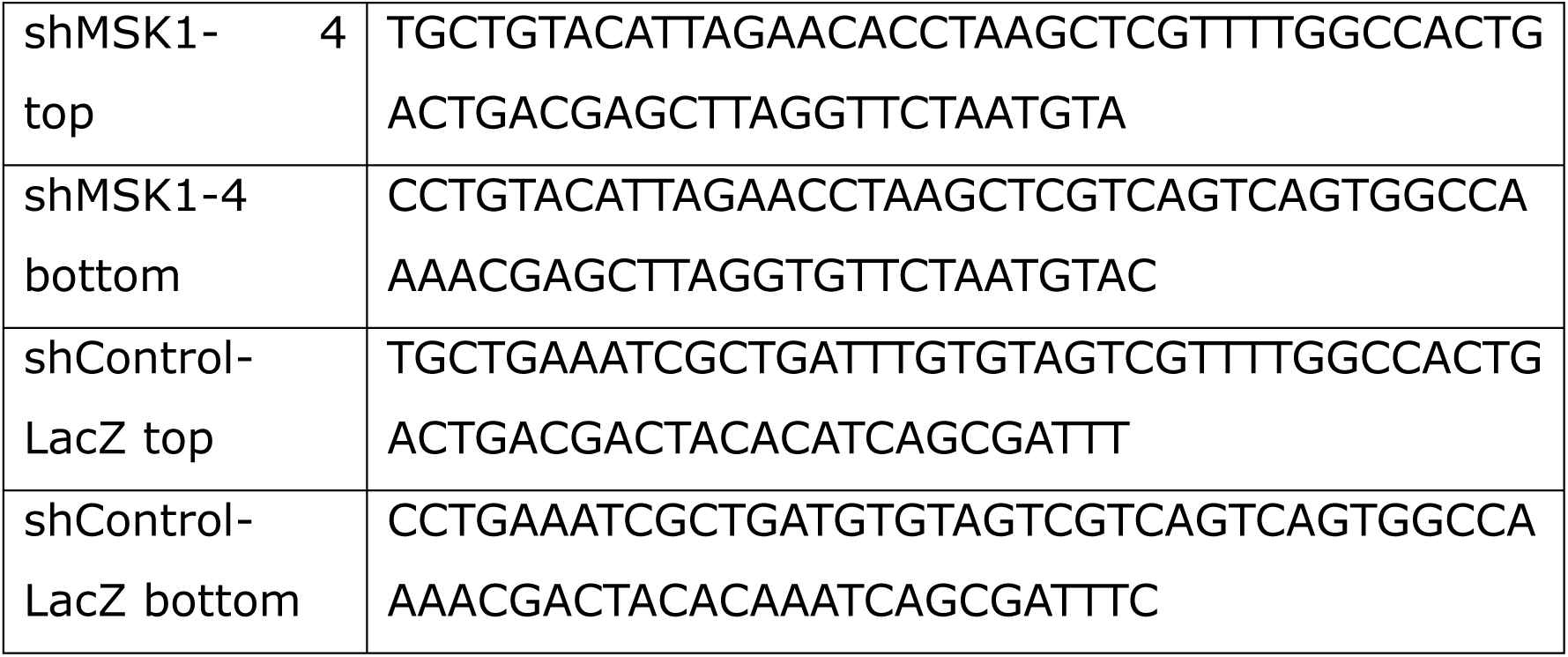
oligonucleotides used to generate the miRNA-based shRNAs.

#### Generation of MSK1 expression vectors

The open reading frame for MSK1 was amplified by PCR from cDNA obtained from C57BL/6J WT mouse brain using the primers MSK1 BamHI-Forward, MSK1 NotI- Reverse, MSK1 NotI-myc-Reverse (Table 5) The PCR product was purified using an agarose gel and co-digested with BamHI and NotI restriction enzymes (New England Biolab) and directly cloned into the pLL-hSYN-hSYN-GFP vector at the BamHI and NotI restriction sites located downstream of the first hSYN promoter. Transformed bacterial colonies were screened by PCR and positive clones were subsequently verified by analysis using restriction enzymes. The MSK1 and MSK1-myc sequences were confirmed by PCR sequencing using specific primers for mouse MSK1 (MSK1-440, MSK1-532, MSK1-955, MSK1-1490, MSK1-1964; Table 5). The MSK1-myc plasmid was mutated to made it resistant to the miRNAs-based shRNAs using the oligonucleotides mut-shM1, mut-shM2, mut-shM3, mut-shM4 (Table 5) and the QuikChange Multi Site-Directed Mutagenesis kit (200514; Agilent; Manufacturer Instructions) in several assays until all the mutations were incorporated. All the PCR products were analysed by Sanger sequencing (primers MSK1-440, MSK1-532, MSK1-955, MSK1-1490, MSK1-1964). The final product was then PCR amplified with the oligos BamH1-Forwad and NotI-myc-Reverse (Table 5) to clone into the pLL-shMSK1 digested with the same restriction enzymes in order to generate the pLL-shMSK1-mutMSK1 vector. The final vector pLL-mutMSK1, containing the coding sequence for the reporter protein DsRED, was generated by amplification of the mutMSK1-myc sequence with the oligos MSK1 NheI Forward and the MSK1 EcoRI-myc-Reverse (Table 5). The pLL-MeCP2-HA expression vector was generated as previously described (Yazdani, Deogracias et al., 2012).

**Table 5:**
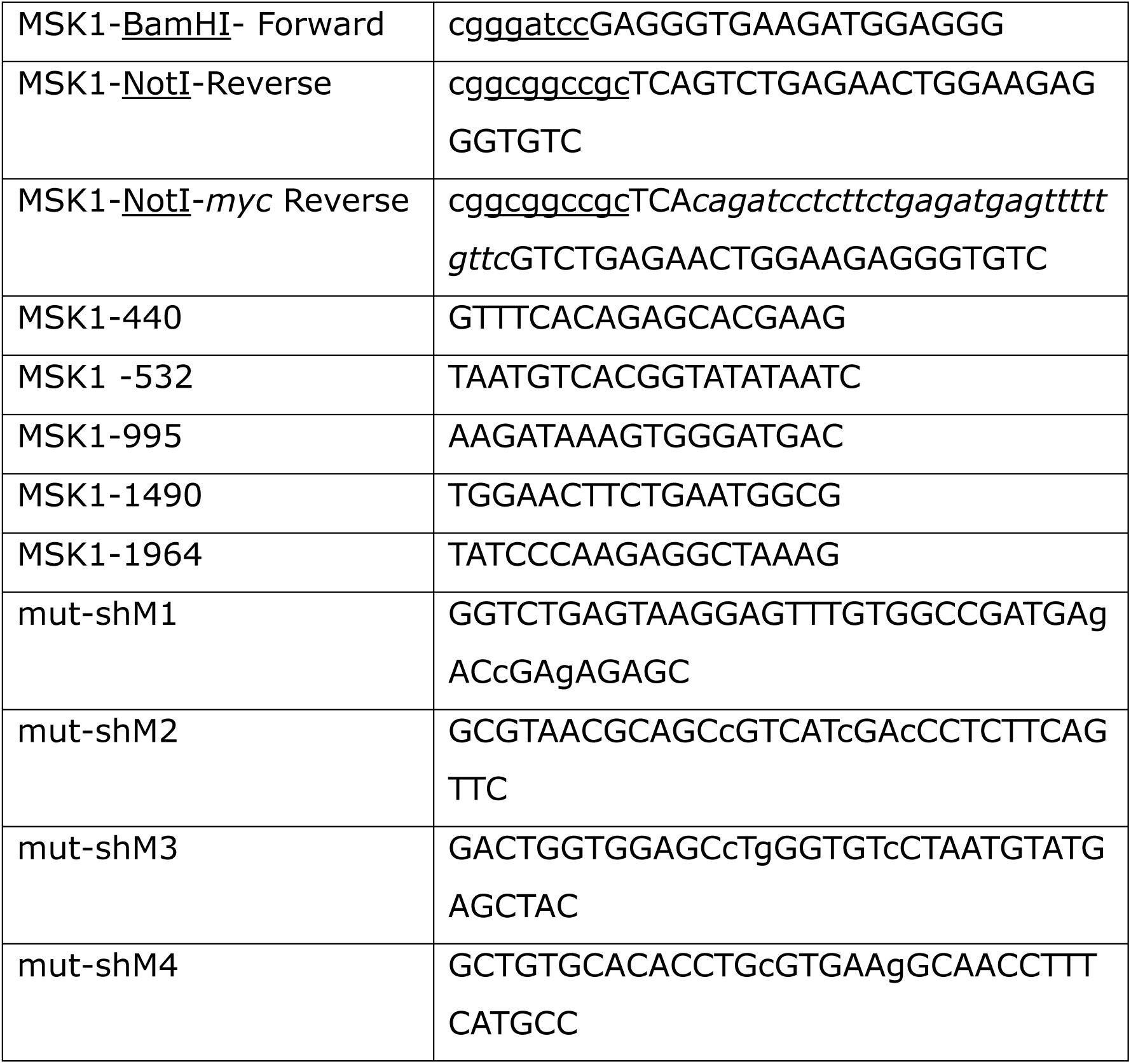
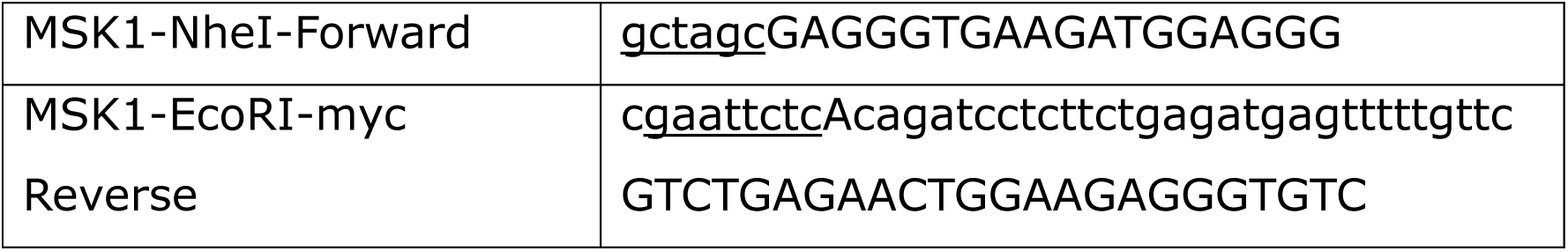
Primers for cloning of MSK1 and Sanger Sequencing and primers to generate the shRNA-resistant form mutMSK1 and mutMSK1myc.

### Cell transfection

Striatal neurons cultured on 24 well plates containing 14 mm diameter coverslips treated with poly-D-lysine and laminin, were transfected 2 days after plating with 1 μg of total DNA with FuGENE 6 transfection reagent (E2691; Promega) at a ratio 1:3.

HEK293-FT cells were plated the day before transfection at a density of 75000 cells per cm^2^. Medium was replaced by fresh DMEM supplemented as before 1h before transfection. For transfection, plasmids were diluted in DMEM and incubated for 20 min at RT with Polyethylenimine (PEI, 23966, Polysciences) at a ratio 1:3 DNA:PEI. 24h after transfection medium was replaced and cells were maintained for 48h after transfection before been collected for obtain total protein lysates or nuclear lysates.

### Immunocytochemistry and neuronal morphology analysis

Striatal neurons were fixed by adding directly to the medium an equal amount of a PBS solution pH 7.4 containing PFA 8% and sucrose 8%. Cells were kept for 20 min at RT and then washed three time with PBS. Blocking solution was applied for 1h (0.1% Triton X-100, 10% normal donkey serum, 5% BSA), and primary antibodies against EGFP (MA1-952; ThermoFisher) diluted 1:200 in antibody solution (0.1% Triton X-100, 10% normal donkey serum, 1% BSA) were applied for 24h at 4°C without shaking. After 3 washes with PBS, donkey-anti-rabbit-Alexa 488 (21206; ThermoFisher) diluted 1:400 in antibody solution containing Hoechst 33258 were applied and kept in the dark at RT for 1h, washed 2 times in PBS and mounted onto glass slides using Mowiol/DABCO mounting media.

Transfected neurons expressing EGFP were visualized using standard epifluorescence under a 5x Neofluar objective. Sholl analyses were performed manually with the help of ImageJ software, setting the origin of the concentric circles in the centre of the soma and separated each 10 μm. A maximum length of 350 μm from the soma was analysed in order to determine the length of the neuronal arborization.

### Purification of nuclear protein lysates

Cells were collected on ice-cold-PBS, centrifuge at 800g for 5 min at 4°C and resuspended in 10 vol. of 250-STM buffer (sucrose 250 mM, Tris-HCl 50 mM pH 8.0, MgCl2 5 mM, NP-40 0.5%, proteases and phosphatases inhibitors). After 10 min incubation on ice, cells were centrifuged at 800g for 10 min at 4°C. The pellet was resuspended in 1 vol. of Nuclear Lysis Buffer (Tris-HCl 20 mM pH 7.5; KCl 300 mM, EDTA 0.2 mM pH 8.0, NP-40 0.1%, glycerol 20% and proteases and phosphatases inhibitors) and incubated for 30 min on ice before been centrifugated at 12000g for 30 min at 4°C. The supernatant, containing the nuclear proteins, was aliquoted and stored at −20°C.

### Co-immunoprecipitation assays

To determine the interaction between MSK1 and MeCP2 independently of the cell context, co-immunoprecipitation was carried using magnetic beads coupled to anti-HA antibodies (SAE0197, Sigma-Aldrich; Manufacturer instructions) on nuclear protein lysates from HEK293-FT cells co-transfected with expression plasmids for MSK1-myc and MeCP2-HA. Immunoprecipitated proteins were thereafter analysed by western blot using monoclonal anti-MSK1 antibody (#3489; Cell Signaling) 1:1000 and polyclonal antibodies anti-MeCP2 (10861-1-AP; Quimigen) 1:500 as control of the immunoprecipitation step.

### Behavioural tests

All the behavioural tests were performed at the Institute of Neurosciences of Castille and Leon (INCyL) animal facility, between 12:00pm and 15:00pm, considering a room with stable temperature, humidity, light intensity and avoided from noises. Before each test, mice were acclimated during 30min in individual homecages.

#### Open field test

Anxiety-like behaviors, exploration and movement were evaluated using this test. Open field arena was made of white fully opaque plexiglas. Square base is 50 cm x 40 cm and walls are 30 cm high to avoide mice to jump over. Mice were placed in the center of the arena and let to free explore during 10 minutes. Measurements as distance travelled (cm), time spent (sec), and number of entries in center or periphery zones were taken and analyzed with tracking software (ANY-maze).

#### Rotarod

Tasks were run with a Rotarod LE 8300 (LSI Letica) to measure motor coordination and balance. Mice were trained for 2 consecutive days before test. Rotarod conditions were setup to increase from 4 rpm to 40 rpm in 5min. Each task was repeated three times, with a 10 min interval between each repetition. Measurements taken were time till fall or time spent on the apparatus (sec) and velocity reached (rpm).

#### Three Chamber Social Test

This task evaluates social approach and interaction with novel inputs. Arena was made of white fully opaque plexiglas and consists of three spaces: middle chamber (where mice are placed to start the test, empty), left chamber (where new and same gender conespecific mouse was situated inside of a cage) and right chamber (where a new object, an orange Lego piece, is placed inside a cage). Mice were allowed to see and sniff freely in the three chambers. In this case, we do not focus on novelty analysis, because both stimuli are new. Our main interest was to assess interaction and exploration differences between conespecific mouse and objects. Habituation was made by placing the mice into the arena with all of the three chambers empty, being them capable of moving freely. Tests start with mouse placed in the empty middle chamber, from where mouse explores and moves between chambers during 10 minutes. Interaction time was measured as the time (sec) the mice spent sniffing around the cages. Trials were recorded and further analyzed using the tracking software *ANY-maze*.

#### Marble Burying Test

This test assesses explorative behaviors and anxiety-like symptoms. Arena consists of homecages filled with 5 cm thick sawdust, with 20 marbles placed on top of it, displayed in 4 rows of 5 marbles each. Each mouse had 30 minutes to freely move and interact with the marbles. We count the number of buryed marbles, considering a marble as buryed if it had 2/3^rds^ covered by the sawdust.

#### Nesting Test

This test evaluates mice capacity to build a place to sleep, hibernate or reproduct. Mice are given a cotton piece (5.1 – 5.2 g) placed in the upper left corner of a homecage filled with 0.5 cm sawdust. Using the parameters described by Lewarch and Hoekstra (2018), score was assigned to the cotton/nest status at 4h and at 24h of starting the test.

#### Forced Swimming Test

Mice were individually forced to swim in a 5L Beaker glass (FB33119, 270mm height, 170mm internal diameter) filled with 2.5L of fresh water (25°C ± 1°C) for 7min. The water was replaced after each trial. Tests were filmed to further analysis of freezing time (sec), a measure of depression-like mice behaviour.The first 2 minutes were not considered to avoid effects caused by the mice stablishment in the Beaker glass.

### Statistical analysis

All statistical analysis were performed using GraphPad Prism 8.0, except the statistical analysis of the densitometry assays that were performed using the IBM SPSS software, version 26. All error bars shown on graphs represent ± standard error of the mean. The statistical tests used in are indicated in the figure legends. Group differences in the number of immunoreactive cells, area and OD were determined using the non-parametric Kruskal-Wallis test.

## Acknowledgments.

We thank Prof. Yves-Alain Barde for discussions and ideas during the initial steps of this study, Prof. Alberto M. Pendás and members of the M.A.M., J.C.A. and R.D. laboratories for stimulating discussions and ideas.

## Author contributions

Conceptualization: R.D.; Investigation: R.D., N.V-A., A.C-L., C.H-C., I.S.F-C., S.G-L., N.M-A.; Methodology: R.D., M.S-M., M.A.M., J.C.A.; Formal Analysis: R.D., N.V-A., A.C-L., C.H-C., I.S.F-C.; Data curation: R.D., M.A.M.; Writing – original draft preparation: R.D., N.V-A., A.C-L., C.H-C., I.S.F-C.; Writing – review and editing: All authors; Supervision: R.D.

---

**Figure EV1.**
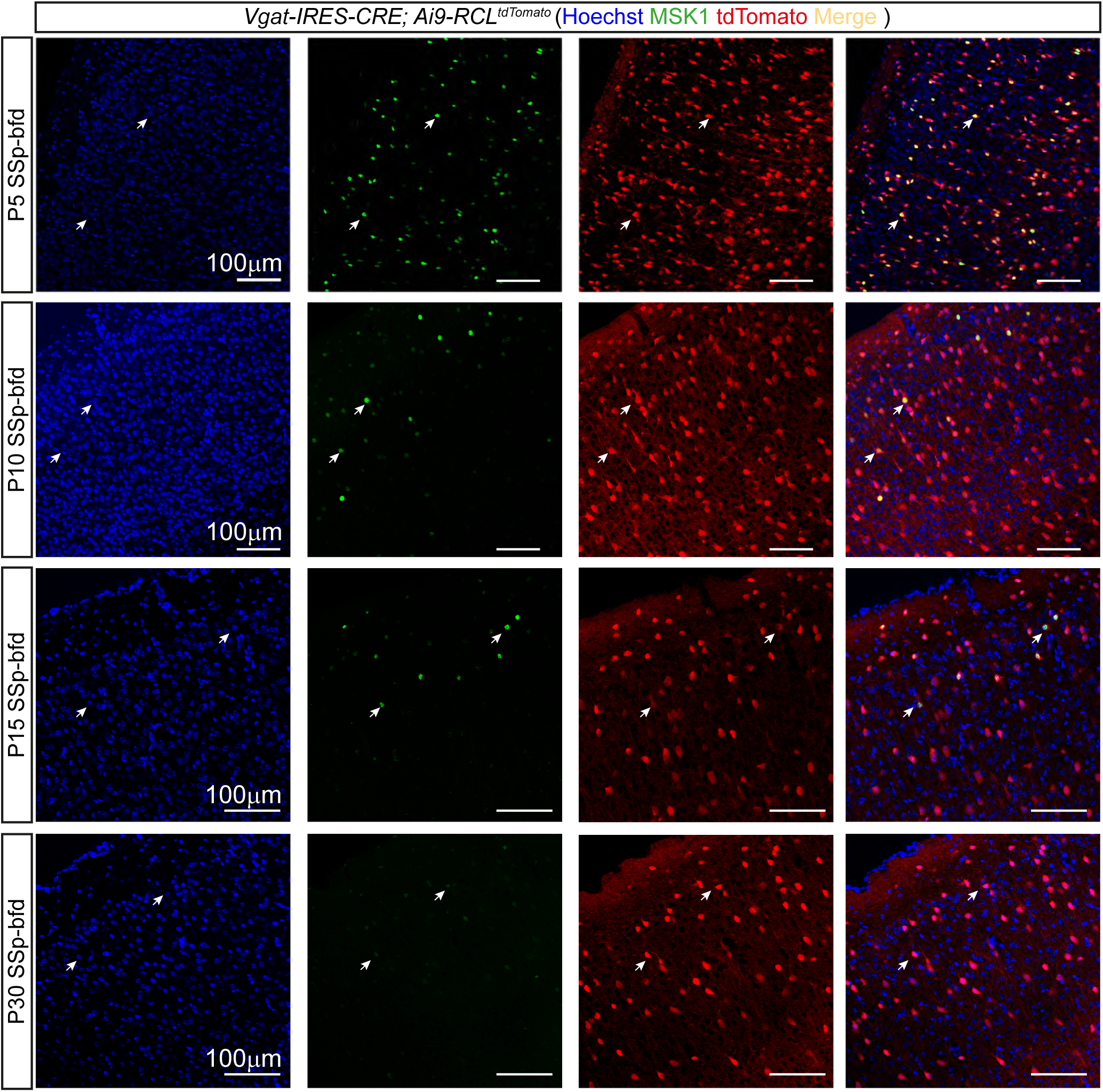
MSK1 expression and distribution pattern change during postnatal brain development. Representative single plane confocal images of P5 to P30 barrel field area of the somatosensory cortex (SSp-bfd) of the *Vgat-IRES-CRE; Ai9-RCL^tdTomato^* mice stained with specific anti-MSK1 antibodies. Endogenous expression of the fluorescent reporter tdTomato labels the soma and neurites of all interneurons in the area. Arrows point to tdTomato^+^ cells that are expressing MSK1.

**Figure EV2.**
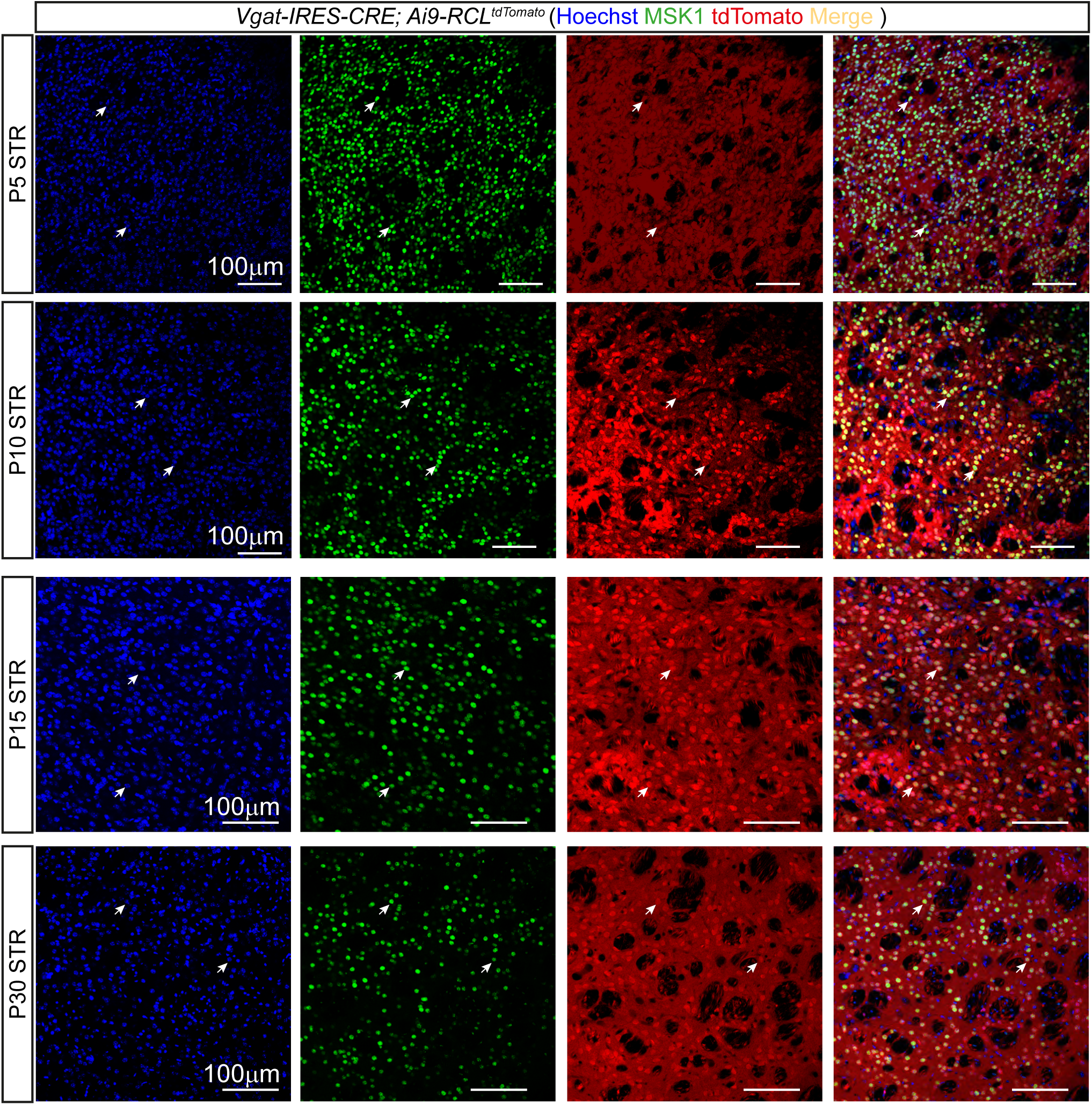
Detection of MSK1 expression pattern in GABAergic neurons in the striatum during mouse brain development. Representative single plane confocal images of MSK1 distribution in the striatum from P5 to P30 in *Vgat-IRES-CRE; Ai9-RCL^tdTomato^* mice expressing the reporter fluorescence protein tdTomato in GABAergic neurons. Arrows point to tdTomato^+^ cells that are expressing MSK1.

**Figure EV3.**
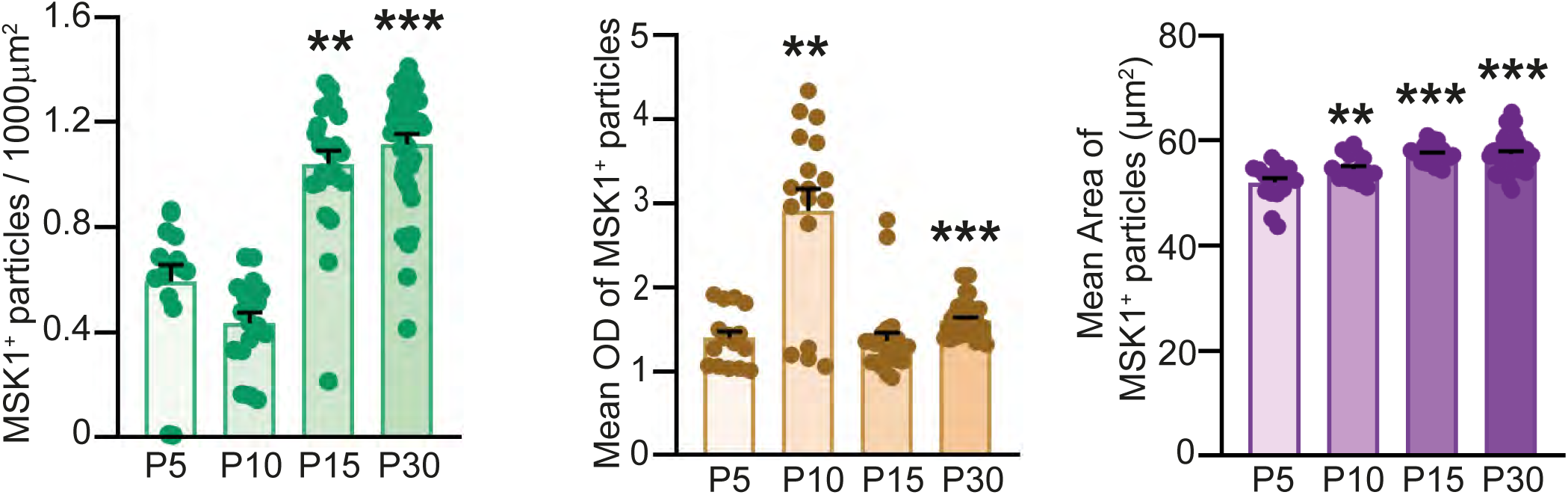
MSK1 expression in striatal neurons development. Statistical analysis of postnatal developmental changes in number, O.D. and size of MSK1^+^ cells in the striatum. (A) Number of MSK1^+^ particles in 1000 µm^2^. (B) Mean optical density (OD) of MSK1 + particles. (C) Mean area of MSK1+ particles (mean ± SEM; n=3. Non-parametric Kruskal-Wallis’ test followed by *post hoc* Bonferroni test; \**P*<0.05, \*\**P*<0.01, \*\*\**P*<0.001).

**Figure EV4.**
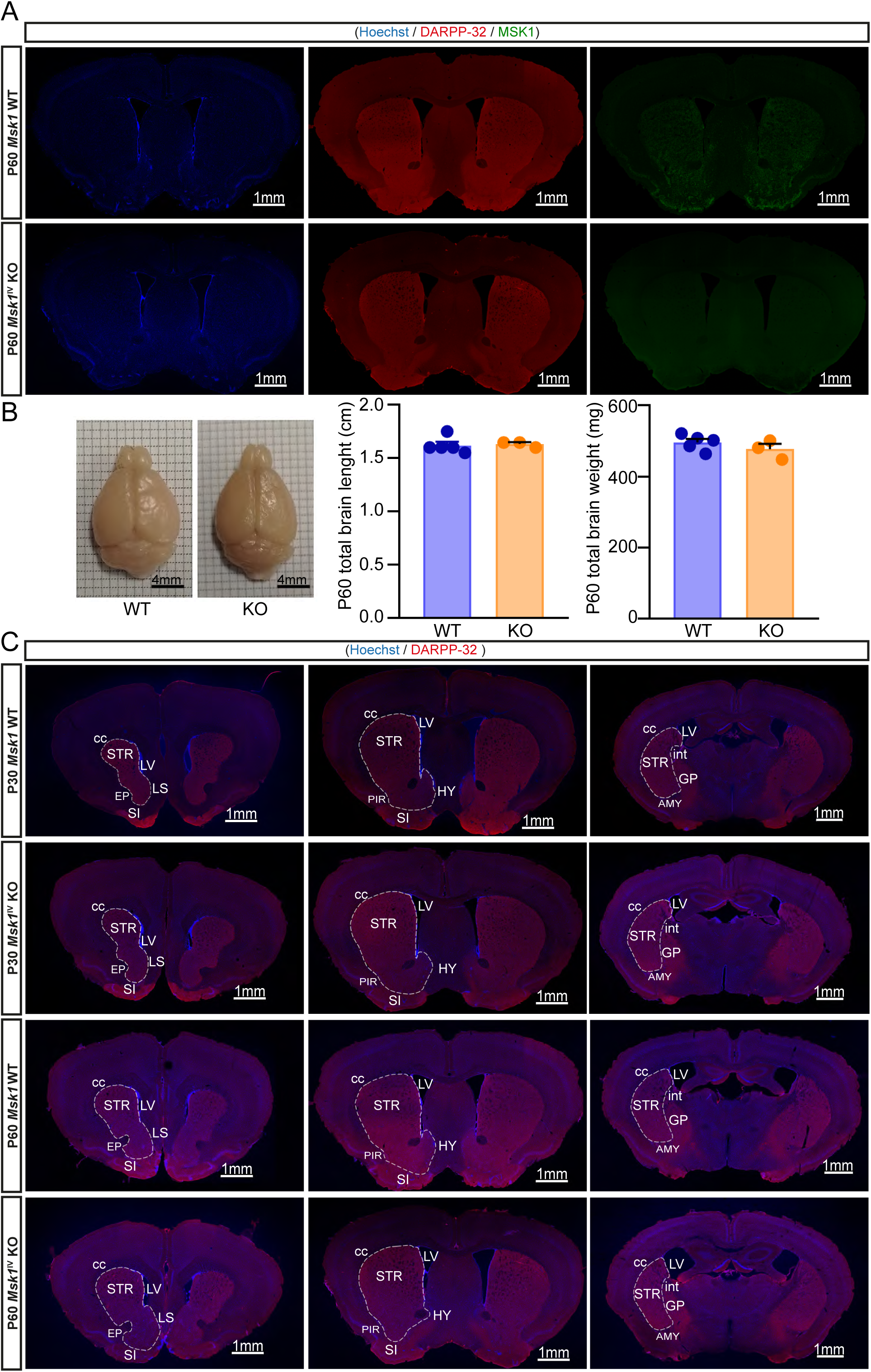
Striatal volume is reduced in the *Msk1^IV^* KO mice. (A) Low power magnification of coronal sections (Bregma ≈ +0.045 mm) from P60 wild type and *Msk1^IV^* KO mice immunostained with anti-MSK1 (green) and anti-DARPP-32 (red) antibodies. (B) Examples of a brain of a wild type and a *Msk1^IV^*KO mice at 2 months old after PFA intracardial perfusion. Brain length and weight measurements from P60 WT and *Msk1^IV^* KO mice.(mean ± SEM; n=5 wild type mice and n=3 *Msk1^IV^* KO mice. two tailed unpaired Student’s t test). (C) Representative images of coronal sections (rostral: Bregma ≈ +1.42 mm, medial: Bregma ≈ +0.045 mm and caudal: Bregma ≈ −1.055 mm) of WT and *Msk1^IV^* KO mice immunostained for DARPP-32 and used for further striatal volume determination applying Cavalierís principle, using the Allen Brain Atlas and considering both the staining and the anatomical boundaries of the striatum. Scale bar in all images is 1mm. STR=striatum, LV=lateral ventricle, LS=lateral septum, SI=substantia innominata, EP=endopiriform cortex, HY=hypothalamus, PIR=piriform cortex, cc=corpus callosum, int=internal capsule, GP=globus palidus, AMY=amygdala.

**Figure EV5.**
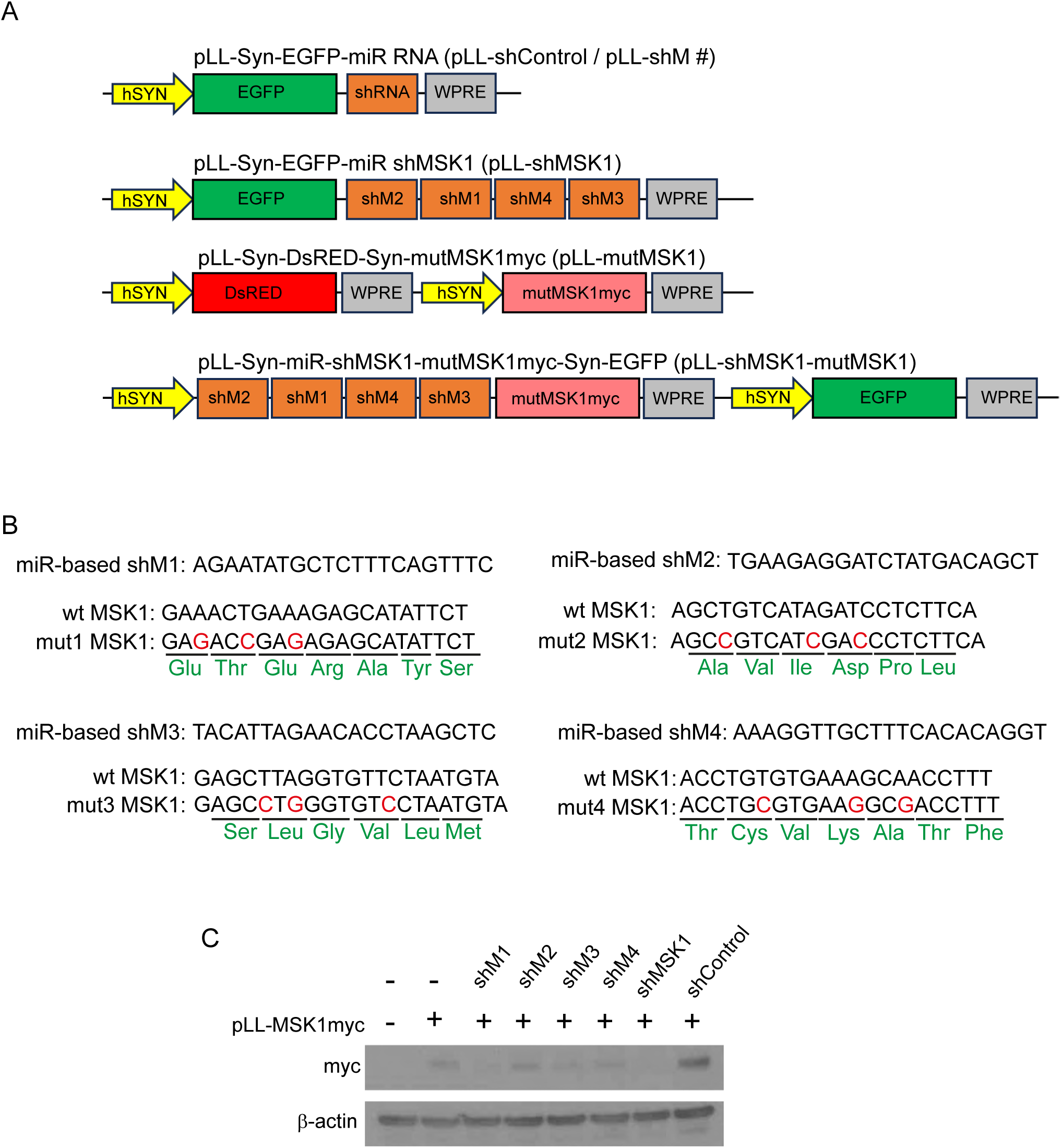
Validation of human synapsin-promoter expression vectors containing miRNA-based shRNAs and a resistant form of MSK1 (mutMSK1). (A) Scheme of the expression vectors for individual miRNA-based shRNAs. Control miRNA-based shRNA (shControl) against β-galactosidase. pLL-shM # (shM1, shM2, shM3 and shM4) are miRNA-based shRNAs against different regions of MSK1. The plasmid pLL-shMSK1 contains a cassette with the four miRNA-based shRNAs against MSK1. The expression vector pLL-mutMSK1 ¡s a dual human synapsin-promoter vector allowing the independent expression of the fluorescent reporter protein DsRED and the protein MSK1 containing point mutations in its nucleotide sequence that confer resistant to the miRNA-based shMSK1-induced degradation. The vector pLL-shMSK1-mutMSK1 allows the expression of the cassette containing the four miRNA-based shRNAs against MSK1, the resistant form of MSK1 and the reporter fluorescent protein EGFP under the control of the promoters human synapsin. (B) Sequences of the miRNA-based shRNAs against MSK1 and the corresponding point mutations that change the nucleotide sequence but not the aminoacidic sequence to mutMSK1. (C) Western Blot showing the efficiency of each miRNA-based shRNA on HEK293-FT cells co-transfected with an expression plasmid for the murine wild-type MSK1 protein containing a myc-tag domain at the end of the 3’CDS.

**Figure EV6.**
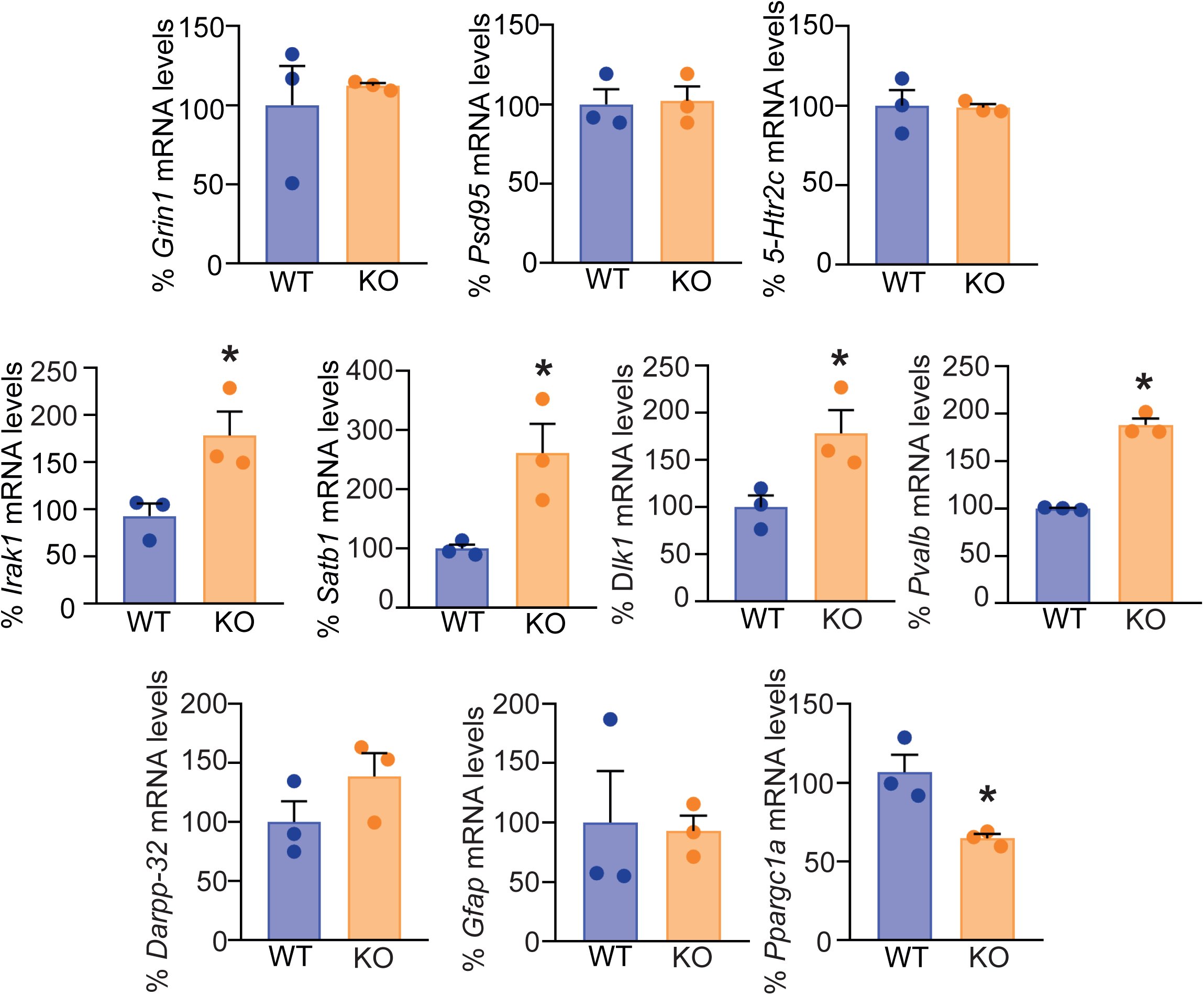
Lack of MSK1 causes gene dysregulation in the striatum. Transcript levels for *Grin1*, *Psd95*, *5-Htr2c*, *Irak1*, *Satb1*, *Dlk1*, *Pvalb*, *Darpp-32*, *Gfap*, *Ppargc1a* analysed by qPCR using specific primers for each. * *P*<0.05. Data were analysed as mean ± SEM, n=3, two tailed unpaired Student’s t test.

